# Circuit and cell-type mechanisms of the spectrolaminar motif of cortical oscillations

**DOI:** 10.64898/2026.11.04.763717

**Authors:** Beatriz Herrera, Shinya Ito, Darrell Haufler, Kael Dai, Ahmed Abdaltawab, Yiti Li, Eli Sennesh, Diego Mendoza-Halliday, André M. Bastos, Anton Arkhipov¹

**Affiliations:** Allen Institute, Seattle, WA; Department of Neuroscience, University of Pittsburgh, Pittsburgh, PA, USA; Department of Psychology, Vanderbilt University, Nashville, TN, USA

## Abstract

Oscillations are a key feature of cortical activity, coordinating spike timing across areas, layers, and cell types. Recent work suggests these rhythms follow a stereotyped organization across cortical depth—the spectrolaminar (SL) motif—with higher-frequency gamma prominent in superficial layers and lower-frequency alpha/beta in deep layers, yet the mechanisms generating it remain poorly understood. Using a bio-realistic model of mouse primary visual cortex, we reproduced the SL motif and found it preserved across diverse visual stimuli. Systematic cell-type– and layer-resolved perturbations showed that the alpha/beta component was intrinsic to every layer, whereas the 50–80 Hz gamma component was predominantly generated within layer 4 (L4) and propagated network-wide through inter-laminar connections. Pvalb interneurons were the principal regulators of gamma, Sst cells shaped low-frequency power, and Vip cells controlled oscillation magnitude. The temporal properties of L4 Pvalb-to-excitatory synapses were especially influential in generating the SL motif. These models and simulation results thus offer a plausible description of circuit mechanisms underlying cortical oscillations and the SL motif and are publicly available to facilitate further studies.

## Introduction

Cortical oscillations coordinate spike timing within and across brain areas, layers, and cell types, supporting functions from sensory processing and selective attention to working memory and learning ^1–6^. These rhythms span multiple frequency bands, each broadly associated with distinct brain states, structures, and putative functions: delta with sleep, theta with cortico-hippocampal communication and spatial navigation, gamma with local cortical computation, etc. Through reciprocal cortical interactions, the thalamus helps generate and pace several of them ^2,3^.

A recently proposed organizing principle suggests that these rhythms are arranged systematically across the cortical depth in a stereotyped pattern, termed the spectrolaminar (SL) motif: higher-frequency gamma is strongest in supragranular and granular layers, while lower-frequency alpha/beta activity (∼8–30 Hz) is strongest in infragranular layers^7^. This laminar frequency gradient has been reported across many neocortical areas in the macaque and, to varying degrees, observed in the marmoset, human, and mouse cortex ^7^. Although its ubiquity has been questioned ^8^, evidence of the SL motif continues to accumulate ^9–13^. The possibility that distinct oscillation types are separated across cortical layers carries far-reaching implications for how computations and dynamics are organized in the cortex: gamma-band activity has been associated primarily with local circuit interactions and feedforward signaling, whereas alpha/beta rhythms have been linked to long-range and feedback-related processing ^14,15^. Together with the SL motif, these observations point to a candidate organizing principle for cortical computation. Importantly, however, the cell-type– and circuit-level mechanisms that give rise to the SL motif remain poorly understood.

Inhibitory interneurons are central to generating cortical rhythms ^2–4,6^, with distinct roles proposed for major subclasses. In mouse primary visual cortex (V1), the spiking of parvalbumin-expressing (Pvalb) and somatostatin-expressing (Sst) interneurons preferentially correlates with gamma- and beta-band oscillations, respectively ^16^. Pvalb cells provide fast perisomatic inhibition and lead the gamma cycle, phase-locking to gamma more strongly than Sst cells ^17^. Additionally, their silencing reduces gamma-band oscillations while increasing low-frequency oscillations ^16^. Sst cells target the dendrites of excitatory cells and also support V1 gamma synchronization—their activation promotes, and their suppression reduces gamma ^16,18^. Vasoactive intestinal peptide-expressing (Vip) interneurons act mainly by disinhibiting Sst cells, locally controlling gamma gain and globally its coherence ^19^. Despite this growing mechanistic picture at the level of individual cell types, it remains unclear how their layer-specific distributions and interactions combine across the cortical laminae and which circuit properties set the dominant oscillation frequency in each layer.

To address these gaps, here we use a biologically realistic, large-scale model of mouse V1 (∼67,000 neurons across layers 1-6 (L1–L6), representing the major excitatory and inhibitory cell types) ^20^. Relative to its predecessor^21^, this model incorporates updated synaptic connectivity derived from large-scale electron-microscopy connectomics ^22^, synaptic dynamics constrained by multi-patch synaptic-physiology measurements ^23^, and, most importantly, is optimized by a deep-learning training procedure (backpropagation through time) that tunes synaptic weights so that the network reproduces cell-type-specific firing statistics and tuning properties measured in vivo with Neuropixels probes ^20^. Importantly, oscillatory activity was never an explicit construction or training target; thus, any emergent rhythm constitutes an independent prediction of the trained circuit. Because the model includes major cortical cell types and all layers, it enables study of integrated operation of the whole V1 circuit across the cortical laminae. This opens up new avenues for exploration of cortical oscillation mechanisms that, until now, have not been extensively modeled, since most modeling studies of cortical oscillations did not resolve multiple layers or specific differences between them ^18,19,24–26^.

By combining drifting-grating and naturalistic visual stimulation with cell-type– and layer-resolved perturbations—namely, interneuron silencing, single-layer isolation, and targeted swapping of parameters between layers—we examine the cell-type and circuit origins of the SL motif. We find that the V1 model reproduces the SL motif across drifting gratings, natural scenes, and movies. Layer 4 drives 50–80 Hz gamma that propagates to layers 2/3 and 5, alongside slower intrinsic oscillations strongest in layer 6. Pvalb cells most strongly modulate gamma, while Sst cells shape lower-frequency power and Vip cells regulate oscillation magnitude, especially in upper layers, without altering the frequency structure. Layer 4 gamma emerges from multiple circuit properties rather than a single cell type or parameter, with Pvalb-to-excitatory synaptic time constants and amplitudes especially influential. Our work establishes a mechanistic framework for understanding the circuit origins of the SL motif across all cortical layers and major cortical cell types. We made the code, models, and simulation results publicly available to support further research in the field.

## Results

### The spectrolaminar oscillatory motif emerges in the bio-realistic model of mouse V1

To investigate cell-type– and layer-specific mechanisms of visually induced cortical oscillations, we used a bio-realistic large-scale model of mouse V1 ^20^, comprising ∼67,000 point-neuron models distributed across all cortical layers (L1–L6), with 400 μm lateral radius, and representing 19 cell types (Fig. 1A)^26^ ^20^. It consists of one excitatory (Exc) and three inhibitory subclasses— parvalbumin (Pvalb)-, somatostatin (Sst)-, and vasoactive intestinal peptide (Vip)-expressing interneurons—per layer from L2/3 to L6, plus a single inhibitory subclass in L1 (L1 Inh). ^27,28,29,21,22^ A training procedure refined individual synaptic weights using backpropagation through time to match in vivo cell-type-specific firing rates and orientation- and direction-selectivity indices (OSI/DSI) from the Allen Institute Neuropixels Visual Coding dataset ^27^, while keeping the distribution of the weights for each cell-type pair consistent with the experimental measurements ^23^ (see Methods). The model is driven by realistic, data-grounded LGN-to-V1 inputs (see Methods). A complete description of the model and optimization process is provided in Ito et al. (2026).^20^

**Figure 1.**
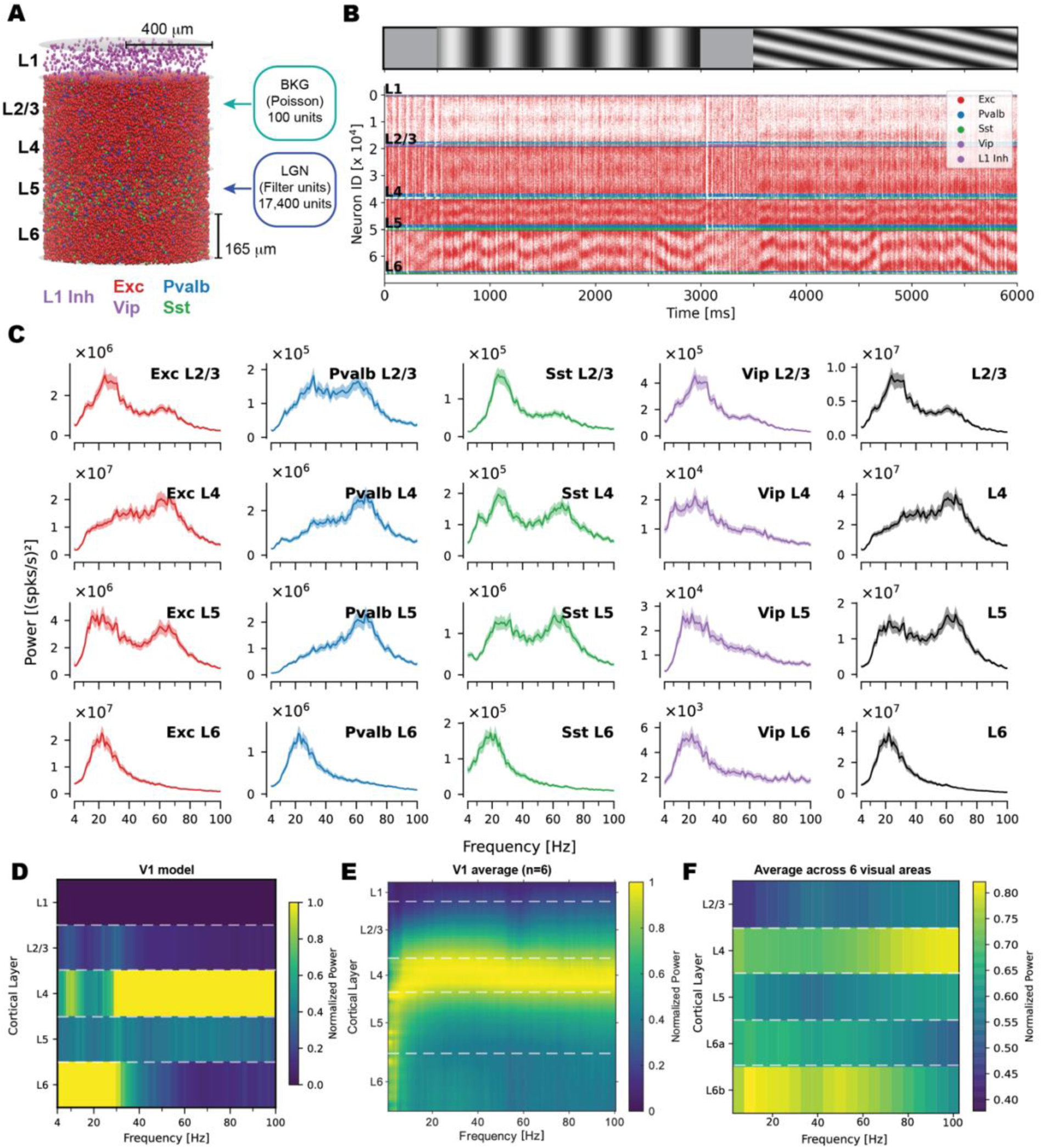
The spectrolaminar motif in the mouse V1 model during drifting-gratings stimulation. **(A)** Schematic of the bio-realistic model of mouse primary visual cortex (V1), displaying excitatory (E) and inhibitory (Pvalb—parvalbumin-; Sst—somatostatin-; Vip—vasoactive intestinal peptide-expressing) cell subclasses distributed across cortical layers L1–L6. Thalamocortical (LGN) and background (BKG) external drives are indicated. The model comprises ∼67,000 neurons that receive input from 100 BKG units and 17,400 LGN filter units. **(B)** Representative spike raster plot of model activity (sorted by cell type and preferred direction) in response to drifting gratings at 0° and 45° directions. **(C)** Cell-type– and layer-specific power spectra of responses to drifting gratings (0.8 contrast, 2 Hz temporal frequency, 8 directions). Each subplot corresponds to a specific cell population and layer, as labeled. Power is expressed in (spikes/s)². Shaded regions here and in all other figures represent ±1.96 times the standard error of the mean (SE) across trials and directions. **(D)** V1 model’s normalized laminar power spectra illustrating the spectrolaminar motif. For normalization, we divided power at each frequency by the maximum power across layers, revealing the characteristic frequency–layer organization: gamma-band dominance in L4 and alpha/beta dominance in L6. **(E)** Experimentally determined spectrolaminar pattern in mouse V1 – mean laminar power spectra across 6 mouse V1 laminar recordings showing LFP power normalized across layers at each frequency (see Methods). **(F)** Experimentally determined spectrolaminar pattern in mouse visual cortex – mean power spectra of each layer. As in E, we divided each trial’s LFP power by its maximum across the probe’s channels at each frequency. We then averaged across the layer’s channels and pooled across trials and probes (83 Neuropixels probes in six visual areas: V1, LM, RL, AL, PM, AM; 14 mice; see Methods).

In these simulations, network activity was recorded as spike trains from all neurons, and cell-type– and layer-specific population firing rates were computed as proxies for laminar oscillatory content (see Methods). This is because point-neuron network simulations output spike times for each neuron in the network but do not produce a biophysical signal such as local field potentials (LFPs), which are typically used in experimental studies to characterize oscillations ^28–34^. While population firing-rate spectra differ from LFP spectra in important ways—most notably by not capturing subthreshold membrane potential fluctuations or dendritic current contributions—they offer a complementary and direct measure of the oscillatory spiking activity for neural populations of interest, not influenced by synaptic and volume-conductance contributions from remote populations, as LFP recordings can be. Ultimately, both measures are useful, but practically, LFP is much more experimentally accessible. We use the V1 model’s firing-rate spectra to understand the phenomenology and mechanisms of oscillations across cortical layers and qualitatively compare our findings with LFP-derived experimental observations. Prior simulation studies using point-neuron models have similarly used population firing-rate power spectra to characterize network oscillations ^19,25,35,36^. We computed power spectra using Welch’s method (see Methods). Raw spectra are reported in units of (spikes/s)^2^, and normalized spectra were derived by dividing power at each frequency by the maximum power across layers at that frequency.

Before characterizing visually evoked dynamics, we studied spontaneously evoked oscillations in the trained V1 model (see Methods). The spontaneous power spectra across layers and cell types were dominated by low frequencies, peaking in the alpha/beta range (Suppl. Fig. 1A), consistent with the predominantly low-frequency structure of spontaneous cortical activity and the stimulus-induced nature of gamma in V1 ^37,38^.

In response to drifting gratings (0.04 cycles/deg spatial frequency, 0.8 contrast, 2 Hz temporal frequency, eight directions, ten trials each), the model generated robust visually evoked spiking activity across all layers and cell types (Fig. 1B). We computed the power spectra at two levels: cell-type-specific spectra, from the population firing rate of each cell type within a layer, and layer-specific spectra, from the aggregated firing rate of all neurons in that layer. Both revealed a consistent spectral pattern across layers: strong oscillations in the alpha/beta and lower gamma range in L6 (∼15-35 Hz) and bimodal spectra in L2/3, L4, and L5, with one component similar to that in L6 and the other spanning the higher gamma range of roughly 50-80 Hz (see Fig. 1C). The 50-80 Hz component was particularly prominent in L4, which also exhibited more intermediate-frequency (e.g., 40 Hz) oscillations. Overall, the L2/3 and L5 spectra were weaker than those in L4 and L6, consistent with the sparse firing of superficial layers ^27,39^, as well as the high neuronal density of L4 and the large neuron count of the thick L6 ^20–22,40^.

To visualize the frequency structure throughout cortical layers, we normalized the laminar power at each frequency by the maximum power across layers (Fig. 1D; Suppl. Fig. 1D). This normalization revealed the SL motif: the normalized alpha/beta power peaked in L6, and normalized gamma power peaked in L4, while intermediate layers occupied intermediate spectral positions. This frequency–depth organization was preserved during spontaneous (gray-screen) activity (Suppl. Fig. 1B). Furthermore, we performed a new analysis of experimental LFP data from the mouse visual cortex, in addition to the previously published SL motifs from the mouse and primate visual cortex ^7^. These included data from laminar probe recordings in mouse V1 (data from ^41^; the mean across 4 mice (6 probes) in Fig. 1E) and an average across several mouse cortical visual areas (data from ^42^; Fig. 1F). While the resulting SL patterns differ in detail, they agree on the main trend: high frequencies are prevalent in L4 and lower frequencies in L6, and the SL motif observed in our simulations (Fig. 1D) is consistent with this observation.

Notably, the L4 gamma component at 50-80 Hz observed in our V1 simulations was not caused by thalamocortical drive since LGN units in our model did not exhibit any 50-80 Hz peaks in their spectra (Suppl. Fig. 1C). While cortical narrowband gamma oscillations at ∼60 Hz, reflecting thalamic oscillations, have been reported in experimental studies ^43–46^, our simplified model of the thalamic drive has no oscillatory activity above ∼5 Hz. Thus, the cortical oscillation structure we report mostly reflects intrinsic properties of the V1 network, rather than oscillatory influences from the thalamus, other subcortical regions, or other cortical areas.

To confirm that the distinct spectral peaks are independent oscillatory modes rather than harmonics of a single fundamental, we decomposed the L4 population firing rate by empirical mode decomposition (EMD ^47^) and tested each pair of intrinsic mode functions for harmonic relationships (three-criterion framework of ^48^; see Methods). No pairing met all three criteria in any of the 80 simulated trials (0/320; Suppl. Fig. 2), indicating that the co-existing components are independently generated oscillatory modes rather than harmonic distortion of a common fundamental.

### The spectrolaminar motif is preserved across diverse visual stimuli

To test whether the SL motif is specific to the periodic temporal structure of drifting gratings or reflects a more general property of the V1 circuitry, we presented the V1 model with naturalistic visual stimuli (see Methods): eight sets of 10 natural images (natural scenes, each shown for 250 ms) and two natural movie clips, each divided into eight 3-second segments (ten trials per segment; Fig. 2A–B). Firing rate distributions were largely consistent across stimulus types (Suppl. Fig. 3). A difference was observed for the Sst cells, which showed reduced firing rates in L2/3-5 and increased in L6 during natural movies, relative to the other stimuli.

**Figure 2.**
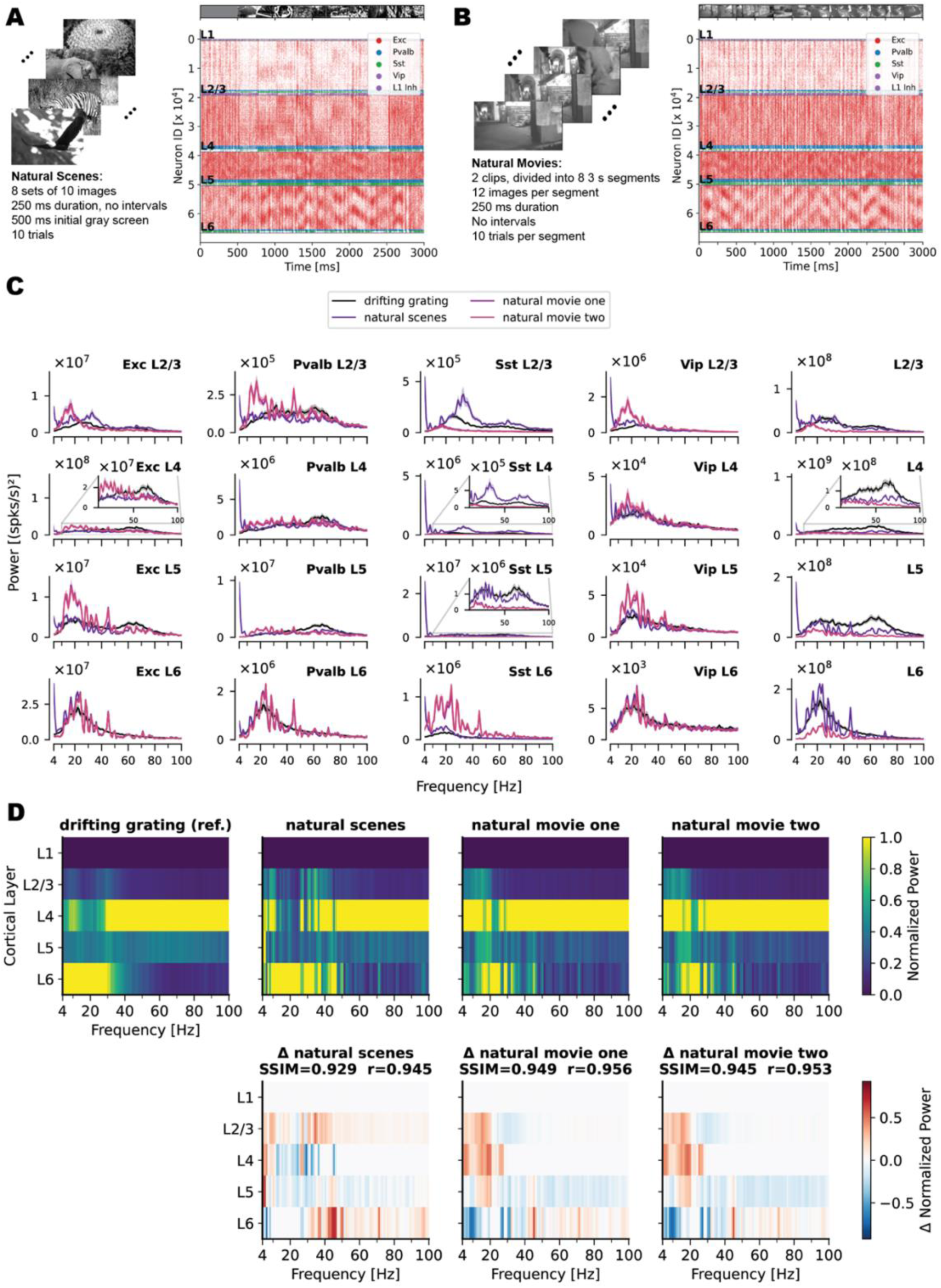
The spectrolaminar motif is preserved across diverse visual stimuli. **(A–B)** Representative spike raster plots (sorted by cell type and preferred orientation) of model activity evoked by natural scenes (A) and natural movies (B). **(C)** Cell-type– and layer-specific power spectra in response to natural scenes, natural movie 1, and natural movie 2. **(D)** Normalized laminar power SL maps for drifting gratings, natural scenes, natural movie 1, and natural movie 2 (top row). Normalization was performed by dividing all values at a given frequency by the maximum power at that frequency across layers. The second row shows the SL map for each naturalistic stimulus minus that for the drifting grating, highlighting deviations from the reference motif. Structural similarity index (SSIM) and Pearson correlation between each naturalistic stimulus and drifting grating SL maps are shown, quantifying preservation of the SL motif across stimulus categories.

Cell-type– and layer-specific power spectra for naturalistic stimuli resembled those observed under drifting gratings (though, for some of them, the overall magnitude was somewhat weaker under naturalistic stimuli relative to drifting gratings). Specifically, the 50-80 Hz gamma component was most prominent in L4, and an alpha/beta component dominated in L6 (Fig. 2C). Laminar SL maps for all three naturalistic stimulus categories were consistent with that for drifting gratings. They showed high structural similarity (SSIM) and Pearson correlation with the drifting grating SL map (Fig. 2D, top row). The difference maps (Fig. 2D, second row) showed only modest deviations, concentrated at lower frequencies, which were stronger across layers in naturalistic stimuli than in drifting gratings.

Because the SL motif persists across stimulus types, we next asked whether it is robust to changes in stimulus parameters. Returning to drifting gratings, we varied stimulus contrast and temporal frequency and found that, although these parameters modulated the magnitude of the oscillatory components, the underlying laminar organization was preserved (Suppl. Fig. 4). Increasing contrast from 0.05 to 0.8 progressively suppressed the 15-35 Hz component in L2/3, L4, and L5, while enhancing 50-80 Hz gamma power, especially in L4 and L5 (Suppl. Fig. 4A). Notably, Sst and Vip population spectra showed opposing contrast modulation: Sst activity increased with contrast while Vip activity decreased (most clearly visible in L2/3, where Vip neurons are most abundant), consistent with mutual inhibition between these two interneuron subclasses and with prior experimental observations of opposing trends in contrast tuning of these cell types ^23,49–51^. Temporal frequency (TF) also modulated oscillations: low TFs (1–4 Hz) suppressed 15-35 Hz oscillations in L2/3, L4, and L5 and promoted 50-80 Hz gamma oscillations in L4 and L5, while high TFs (≥8 Hz) suppressed the 50-80 Hz gamma rhythm (Suppl. Fig. 4B). Remarkably, L6 oscillations in the alpha/beta and low-gamma range were largely unaffected by contrast and TF, except for the amplitude modulation in that range of the L6 Sst neurons, indicating inter-layer influences. Across all contrasts and temporal frequencies, the characteristic laminar arrangement of the SL motif (gamma concentrated in L4, alpha/beta dominant in L6) was maintained.

### Inhibitory interneuron subclasses differentially regulate oscillatory power and frequency across layers

To assess each inhibitory interneuron subclass’s contribution to the SL motif, we silenced Pvalb, Sst, or Vip cells with hyperpolarizing current throughout the simulation (Methods) and compared laminar power spectra with baseline (Fig. 3). Representative rasters show responses to Pvalb and Sst silencing (Fig. 3A–B). Silencing either subclass most strongly altered cell-type firing rates, increasing firing throughout the network (Suppl. Fig. 5A). Vip silencing left firing-rate distributions similar to baseline (Suppl. Fig. 5A).

**Figure 3.**
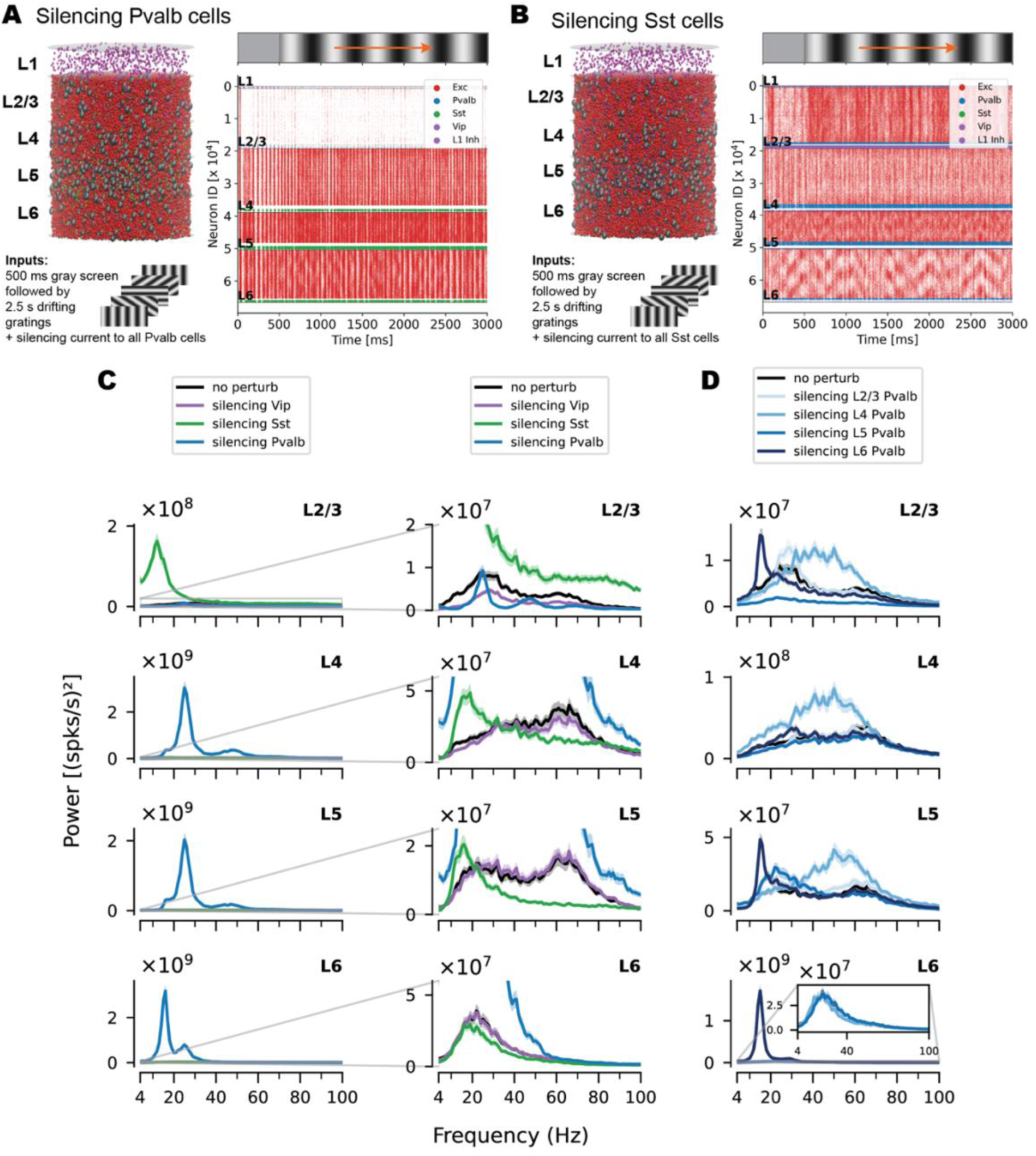
Inhibitory interneuron subclasses differentially regulate oscillatory power and frequency across layers. **(A–B)** Representative spike raster plots (0° direction, single trial) showing V1 network activity during selective silencing of all Pvalb cells (A) and all Sst cells (B). Silenced populations are indicated by enlarged gray spheres (A: Pvalb; B: Sst) in the V1 model. **(C)** Laminar power spectra following whole-population silencing of Vip, Sst, or Pvalb cells, compared to the unperturbed baseline (“no perturb”). **(D)** Magnified view of panel C, highlighting layer-specific frequency shifts. **(E)** Laminar power spectra following layer-by-layer Pvalb silencing (color-coded), showing the contribution of each layer’s Pvalb population to the network-wide oscillatory pattern.

Silencing each of the three interneuron subclasses produced distinct and complementary effects on the SL motif. Silencing all Vip cells reduced the overall magnitude of oscillatory power across layers without substantially altering dominant frequencies or the cross-layer spectral structure (Fig. 3C; Suppl. Fig. 5A). This was particularly noticeable in L2/3, where Vip density is highest ^20,22,52^— supporting a disinhibitory gain-control role in which Vip, by inhibiting Sst, amplifies ongoing oscillations without setting their frequency ^19^. Silencing all Sst cells increased alpha/beta power (10-20 Hz) across all layers (Fig. 3C; Suppl. Fig. 5A). Silencing Pvalb cells had the strongest effect (Fig. 3C; Suppl. Fig. 5A), sharply increasing excitatory firing in L4–6 (Sst silencing acted more in L2/3) and abolishing the 50–80 Hz gamma network-wide, shifting all layers to a bimodal spectrum: L2/3–5 peaked at ∼25 and ∼45 Hz, L6 at ∼15 and ∼25 Hz. This shift in the laminar oscillatory gradient toward alpha/beta suggests that Pvalb cells facilitate and regulate 50-80 Hz gamma, whereas Sst cells more strongly influence alpha/beta and lower gamma frequencies.

The V1 model resolves diverse cell types and all cortical layers, enabling us to investigate how inhibitory cells in specific layers shape oscillations. Silencing Pvalb cells layer by layer produced distinct, pronounced effects (Fig. 3D; Suppl. Fig. 5B). Silencing L4 Pvalb cells promoted broadband 50-80 Hz gamma oscillations across the network (except in L6), confirming the key role of L4 Pvalb cells in controlling the network-wide gamma activity. Silencing L6 Pvalb cells selectively increased ∼15 Hz power in L2/3 and L5-6, with a slight suppression of 50-80 Hz gamma in L2/3 and L5 and no detectable effect in L4, suggesting a modulatory role for L6 Pvalb cells. Silencing Pvalb cells in L2/3 or L5 had substantially smaller effects.

Layer-specific Vip and Sst silencing produced more localized effects (Suppl. Fig. 6; firing rate distributions shown in Suppl. Fig. 5C-D). Silencing L2/3 Vip cells reduced oscillation magnitude without altering the spectral structure (Suppl. Fig. 6A), consistent with their gain-control function^19^. Silencing L5 Sst cells increased 10–30 Hz power in L2/3 and substantially decreased 50-80 Hz gamma power in L5 (Suppl. Fig. 6B), pointing to an L5-specific Sst-dependent mechanism that promotes gamma oscillations locally. Silencing Sst or Vip in other individual layers had minor effects.

### Layer 4 is the predominant source of the 50-80 Hz gamma oscillations across the V1 network

The V1 network intrinsically generates two oscillatory components: 15–35 Hz alpha/beta/low-gamma and 50-80 Hz high-gamma. Their expression varies across layers and reflects layer-specific cell populations. We next investigated which circuit elements generate each. To address this, we performed two complementary perturbations: silencing and isolated-layer simulations. In the silencing perturbation, we silenced all cells in a layer to test whether the remaining network could sustain the laminar oscillation gradient. In the isolated-layer perturbation, we silenced every layer except one (e.g., left L4 active) to test whether its intrinsic circuitry could generate the oscillatory pattern without input from other layers.

Fig. 4A-B shows representative raster plots of network responses to L4 perturbations (cell-type-specific firing-rate distributions: Suppl. Fig. 7A–B). Fig. 4C-D shows silencing and isolated-layer perturbation results for each layer. Silencing L1, L2/3, or L6 had minimal effects beyond the silenced layer. Silencing L4 most strongly affected the network, suppressing 50-80 Hz gamma oscillations and promoting lower frequencies. Silencing L5 caused minor spectral changes in L4 and L6 without considerably altering the key frequencies. Isolated-layer perturbations further emphasized L4’s distinct role (Fig. 4D). In isolation, L2/3 and L6 showed little or no 50-80 Hz activity, oscillating mainly at 15-50 Hz. Isolated L5 retained a reduced 50-80 Hz component, while isolated L4 sustained gamma oscillations alongside lower-frequency activity. Its spectrum remained largely unchanged from the unperturbed V1 network.

**Figure 4.**
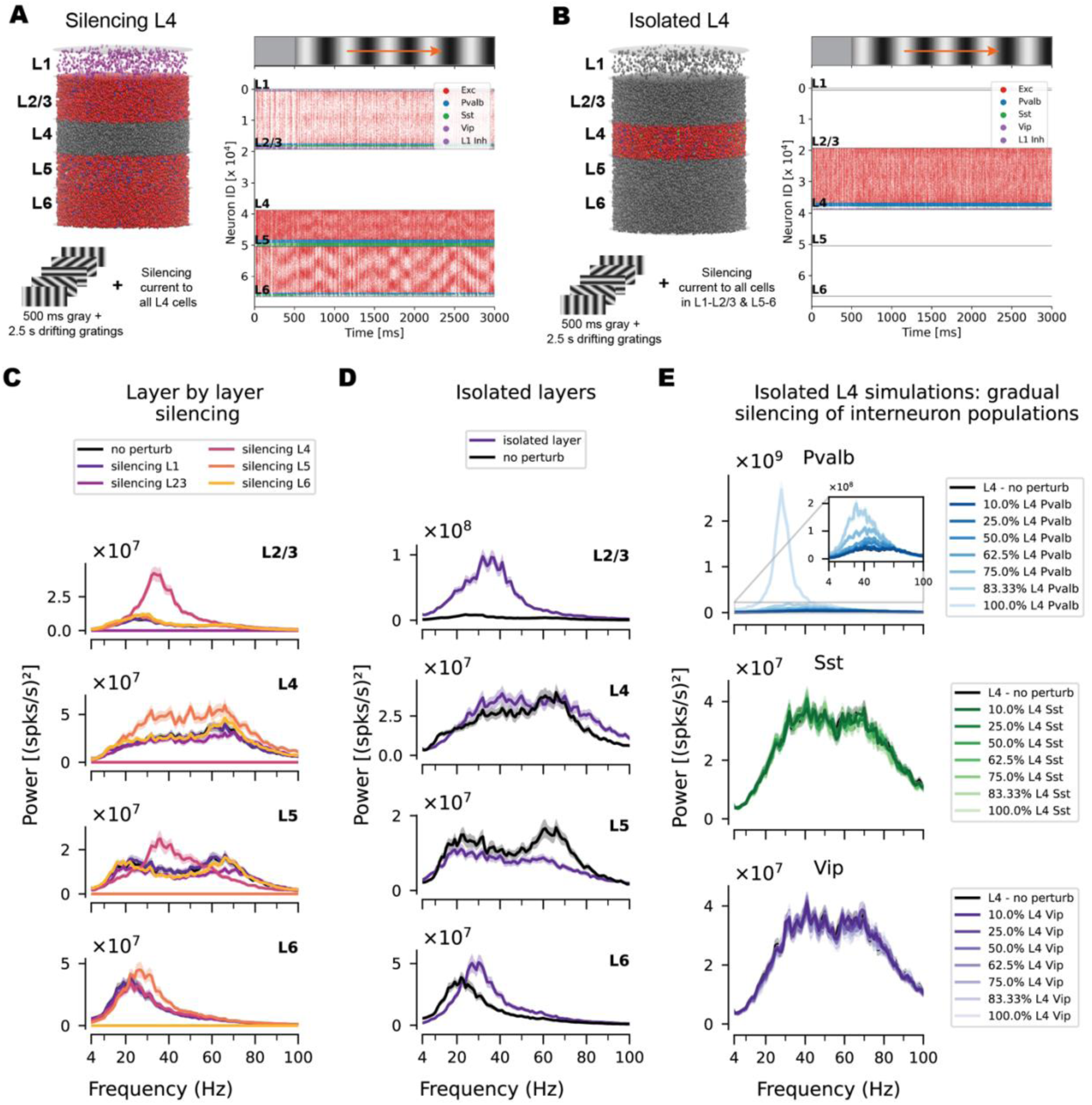
Layer 4 is the predominant source of the 50-80 Hz gamma oscillations across the V1 network. **(A)** Representative spike raster plot (0° direction, single trial) during selective silencing of all L4 cells (gray spheres in the image of the V1 model). **(B)** Representative raster plot from an isolated L4 simulation, in which all cells in L1, L2/3, L5, and L6 were silenced (gray spheres) while L4 remained active. **(C)** Laminar power spectra following layer-by-layer silencing of all cell populations (one layer at a time, with the other layers remaining active). **(D)** Power spectra from simulations of isolated layers. **(E)** L4 power spectra following silencing of different fractions of neurons within inhibitory subclasses in L4 (Vip, Sst, or Pvalb).

To investigate the potential contributions of inhibitory cell types to L4’s special role, we silenced different fractions of these cell types in isolated L4 simulations (Fig. 4E; cell-type-specific firing-rate distributions: Suppl. Fig. 7C-E). These simulations confirmed that, in our model, Pvalb cells are the dominant regulators of L4 50-80 Hz gamma, consistent with the network-wide interneuron silencing results (Fig. 3C). Sst silencing slightly reduced the L4 50-80 Hz gamma amplitude, while Vip silencing had minimal impact on spectral frequency (Fig. 4E).

### Temporal properties of Pvalb-to-excitatory synapses govern Layer 4 50-80 Hz gamma oscillations

After identifying L4 as the primary source of 50-80 Hz gamma oscillations, we examined which circuit features determine its specific oscillatory frequency. PING and ING models suggest inhibitory decay sets gamma oscillation periods: faster decay lets excitatory neurons recover sooner, supporting higher frequencies ^53–56^. Therefore, L4 gamma oscillations likely depend on synaptic-current temporal properties (time constants, amplitudes, and latencies), which we derived in our model from the Allen Institute multipatch physiology dataset ^23^. Examination of these parameters (Fig. 5A, Suppl. Fig. 8) showed that L4 recurrent connections—especially Pvalb-mediated inhibitory-to-excitatory (I-to-E) connections—have faster kinetics, with shorter slow-component time constants than corresponding connections in other layers, consistent with PING and ING predictions. Fig. 5A highlights L4 and L6 to illustrate these differences, as L6 exhibits robust lower-frequency oscillations but little to no 50-80 Hz gamma (Fig. 1C). Consequently, we tested whether layer-specific synaptic temporal properties directly influence the L4 gamma frequency.

**Figure 5.**
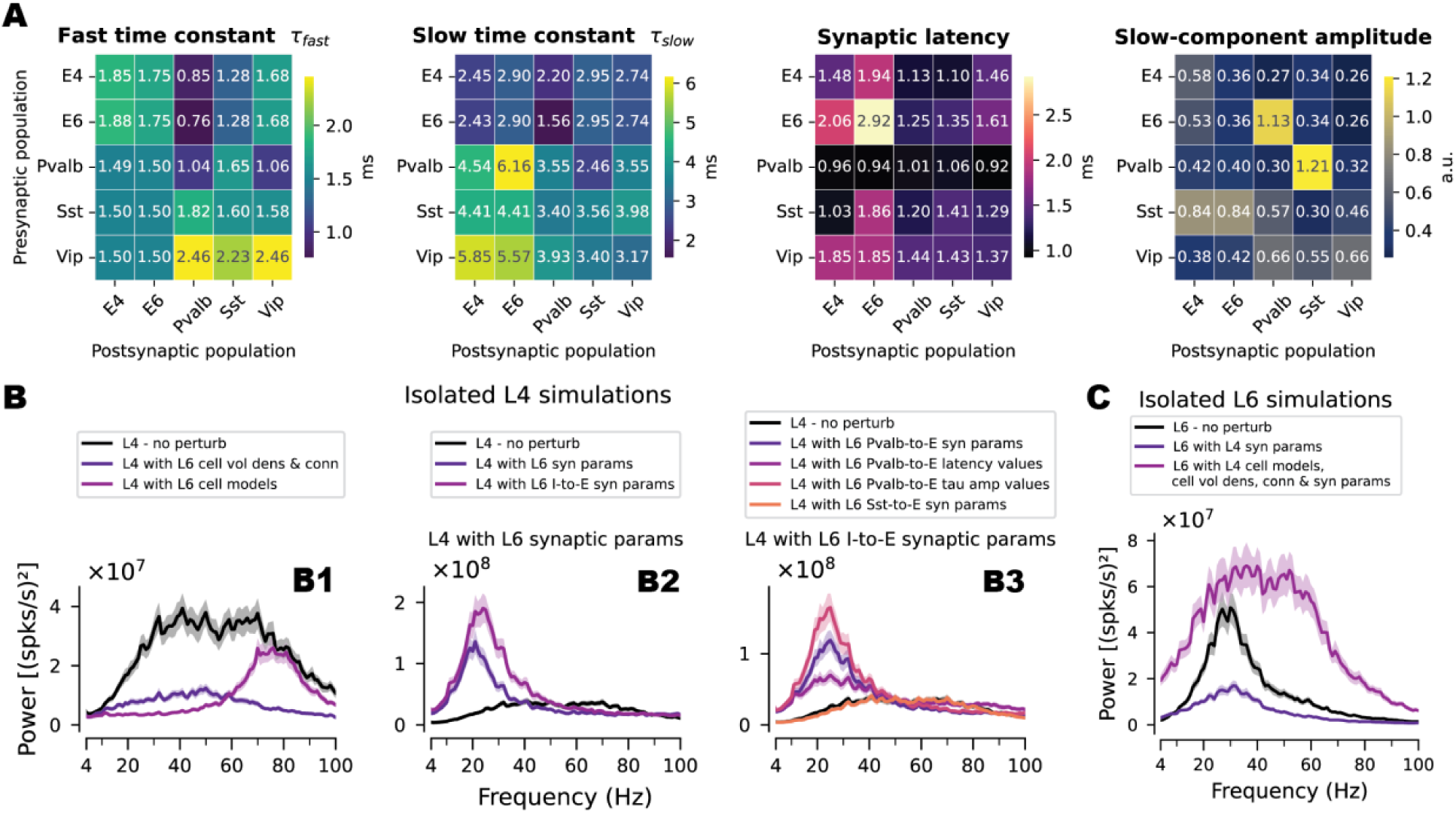
Synaptic parameters of intra-layer recurrent connectivity in L4, especially from the Pvalb population, control the 50-80 Hz gamma oscillations. **(A)** Subset of V1 model time constants (taken from experimental data ^23^) for the fast (1st panel) and slow (2nd panel) components of the double-alpha synaptic currents; the synaptic time latency (3rd panel) and the amplitude of the slow component of the double-alpha synaptic currents (4th panel). The plots highlight L4 and L6 excitatory-to-excitatory and excitatory-to-inhibitory connections within and between these layers, with the y-axis indicating the presynaptic population and the x-axis the postsynaptic population. **(B)** Power spectra from simulations of the isolated L4 network, comparing the unperturbed baseline (“L4—no perturb”) with model variants that systematically replaced L4 network parameters with those from L6. **A1:** Variants swapping cell volume density and within-layer connectivity (“L4 with L6 cell vol dens & conn”) or intrinsic electrophysiological parameters of cell models (“L4 with L6 cell models”). **A2:** Variants swapping all synaptic parameters (“L4 with L6 syn params”) or only inhibitory-to-excitatory (I-to-E) synaptic parameters (“L4 with L6 I-to-E syn params”). **A3:** Targeted variants replacing Pvalb-to-excitatory synaptic parameters—all parameters (“L4 with L6 Pvalb-to-E syn params”), only synaptic latency (“L4 with L6 Pvalb-to-E latency values”), or only time constants and alpha-function amplitude (“L4 with L6 Pvalb-to-E tau amp values”)—and the Sst-to-excitatory counterpart (“L4 with L6 Sst-to-E syn params”). **(C)** Power spectra for simulations of the isolated L6 network, comparing the unperturbed L6 baseline with variants in which L6 network parameters were replaced with those from L4: L6 using L4 synaptic parameters alone (“L6 with L4 syn params”) and L6 using all internal parameters from L4 (i.e., cell models, cell volume density, connectivity, and synaptic parameters are borrowed from L4; the volume occupied by the network and external inputs remain the same as in the original L6 network).

To that end, we constructed a series of isolated L4 model variants in which specific L4 circuit parameters were replaced with the corresponding L6 parameter values and examined how each substitution altered the L4 power spectrum (Fig. 5B). Because even small changes in such circuit parameters can strongly affect the magnitude of spiking activity and network stability, each L4 circuit variant with these changes was retrained to match target in vivo firing rates before spectral analysis (see Methods).

Replacing L4 cell-volume density and within-layer connectivity with L6 values reduced L4 power, shifted the spectral peak to ∼50 Hz, and eliminated L4’s distinct low-gamma (∼40 Hz) and 50-80 Hz gamma (∼65 Hz) peaks (Fig. 5B1; Suppl. Fig. 9E; firing-rate distributions in Suppl. Fig. 9A). Replacing L4 neuron models with L6 models eliminated alpha/beta and low-gamma components and shifted the spectrum to ∼75 Hz (Fig. 5B1; Suppl. Fig. 9A). Thus, intrinsic cell properties shape L4’s low-frequency oscillations but do not primarily determine its 50-80 Hz gamma frequency.

Replacing all L4 synaptic parameters with those from L6 effectively suppressed 50-80 Hz gamma and promoted alpha/beta oscillations (Fig. 5B2; firing rate distributions in Suppl. Fig. 9B), identifying synaptic properties as a dominant factor governing the L4 50-80 Hz gamma. Replacing only I-to-E synaptic parameters produced the same effect. Replacing only the excitatory-to-excitatory (E-to-E) or excitatory-to-inhibitory (E-to-I) synaptic parameters also facilitated alpha/beta and low-gamma oscillations but moderately decreased the 50-80 Hz gamma (Suppl. Fig. 9B, F). These observations suggest that synaptic parameters from multiple cell types shape L4 50-80 Hz oscillations, particularly I-to-E connections.

To identify which I-to-E connections most strongly shaped 50-80 Hz gamma, we replaced each Pvalb-to-E, Sst-to-E, or Vip-to-E synaptic parameter set in L4 with its L6 counterpart, either entirely or only for temporal parameters (fast/slow time constants, slow-component amplitude, and latency; Fig. 5B3; Suppl. Fig. 9C). Replacing only Pvalb-to-E parameters reproduced the suppression seen with a full I-to-E swap (Fig. 5B3). Swapping time constants and amplitudes alone reproduced most of it, whereas latency had a smaller effect. Replacing Sst-to-E parameters had only minor effects (Fig. 5B3; Suppl. Fig. 9G), confirming the primacy of Pvalb-to-E temporal parameters. Moreover, replacing the electrophysiological, synaptic, and connectivity properties of L4 Pvalb cells with those of L4 Sst cells abolished L4 oscillations (Suppl. Fig. 9D, H), whereas the reverse swap left the spectrum largely unchanged. Together, these observations support our earlier conclusion: the alpha/beta/lower-gamma spectrum is a stable property of L4 (and other layers), while 50–80 Hz gamma depends on a narrower set of parameters, especially Pvalb-to-E synaptic kinetics.

To test whether these observations extend beyond L4, we performed the opposite swaps, starting from L6—the layer most resistant to 50–80 Hz gamma—and replacing its parameters with L4 values. Replacing only L6 Pvalb-to-E synaptic parameters (Fig. 5C; Suppl. Fig. 9J) was insufficient to generate 50-80 Hz gamma, reducing only the beta peak magnitude. This likely reflects the weaker LGN input and lower firing rates of isolated L6 (Suppl. Fig. 9I). On the other hand, additionally replacing L6 cell models, cell density, connectivity, and all synaptic parameters with L4 values produced 50–80 Hz gamma in L6 (Fig. 5C; Suppl. Fig. 9J). This modified network still differs from L4: it receives a different LGN drive and has more cells, as L6 is thicker. Firing rates of L6 Exc and Sst cells rose closer to L4’s (Suppl. Fig. 9A, I), whereas L6 Pvalb cells fired considerably less, which may explain the weaker induced gamma relative to L4.

In summary, 50-80 Hz gamma and, consequently, the SL motif are strongly influenced by specific L4 network parameters, particularly the temporal properties of Pvalb-to-E synapses. However, no single parameter or cell type drives them exclusively; rather, multiple influences converge.

## Discussion

The stereotyped organization of cortical oscillations across depth—gamma in granular/supragranular layers and alpha/beta in infragranular layers, collectively termed the SL motif—has been described across species and proposed to underlie the directional flow of information through cortical circuits ^7,14^. However, the cell types and circuit mechanisms underlying this laminar frequency architecture remain unresolved, partly because experimental recordings cannot simultaneously disentangle the contributions of all individual layers and/or cell types. Additionally, prior computational models have usually been limited in their ability to describe multiple interacting cortical layers and more than a few cell types.

In this study, we used a bio-realistic, large-scale model of mouse V1, combined with systematic cell-type– and layer-resolved perturbations, to better understand how visually induced cortical oscillations and the SL motif arise from cell-type-specific circuit properties across the full cortical depth. Classic theoretical work established the canonical excitatory–inhibitory motifs underlying gamma generation (e.g., PING and ING) but did so in comparatively simple network models that did not resolve separate layers or diverse cell types ^53–59^. More recent spiking-network and mean-field studies have resolved multiple interneuron subclasses and their causal contributions to gamma, yet they have done so within non-laminar populations ^18,19,24–26,60^. The published models that do span multiple cortical layers have pursued different questions: multi-layered point-neuron models of V1 characterized population-rate oscillations and their anatomical origins without resolving inhibitory subtypes ^61^; the biophysically detailed, all-layer model of mouse primary motor cortex reproduced layer- and cell-type-specific firing and LFP across behavioral states but did not dissect the cell-type origins of laminar oscillatory frequency structure ^62,63^; and biophysically detailed models of mouse V1 reproduced overall visual responses properties of neural firing ^21^, as well as the laminar LFP, current sinks, and sources across all layers without specifically examining oscillations ^64,65^. Most directly comparable, the multi-layer V1 column model of ^35^ incorporated pyramidal neurons together with Pvalb, Sst, and Vip interneurons across cortical layers, and was fit to reproduce spontaneous and stimulus-evoked cell-type-specific firing rates. In that model, however, rhythmic activity was absent under the tested baseline and stimulus-driven conditions, and fast oscillations emerged only after introducing a spike-timing-dependent plasticity rule that modified excitatory synaptic weights.

Our V1 model represents diverse cell types and all cortical layers, combining the latest neuronal electrophysiology, electron-microscopy connectomics, and multi-patch synaptic data. Trained to reproduce neural activity metrics from in vivo Neuropixels recordings, it receives realistic, data-grounded LGN inputs approximating an animal’s sensory experience during experiments. The model produced visually induced oscillations without additional parameter changes, even though the training did not include oscillations as a target. The observed oscillations robustly formed the SL motif (Fig. 1D), consistent with the observed LFP-derived experimental SL motifs (Fig. 1E-F; Suppl. Fig. 1E). This suggests that the SL motif is determined by factors such as the experimentally constrained feedforward drive, recurrent connectivity, and synaptic physiology.

### The SL motif is a circuit-level property of V1 revealed by visual drive

The model generated two main oscillatory components across visual stimuli, each distributed distinctly across layers (Fig. 1C–D; Suppl. Fig. 1D): a 15–35 Hz alpha/beta (and low-gamma) component present in all layers, and a 50–80 Hz higher-gamma component present in L2/3-5. This frequency–depth organization is the signature of the SL motif, and it persisted across drifting gratings, natural scenes, and natural movies (Fig. 2). Its robustness to stimulus statistics indicates that the SL motif is a fundamental, circuit-level property of V1, revealed by visual input rather than inherited from the stimulus’s temporal structure. These results are consistent with the reported ubiquity of the motif across cortical areas, regardless of their response features ^7^.

Additionally, the model’s normalized SL map (Fig. 1D) reproduced the key features of the experimental maps from mouse V1 (Fig. 1E) and averaged across six visual cortical areas (Fig. 1F): higher-frequency (gamma) power was concentrated in the granular/supragranular layers— peaking in L4 in the model and across L2/3–L4 in the recordings—whereas lower-frequency alpha/beta power was relatively stronger in the deep layers, most evident in L6 (model) and L6b (data). Notably, although the model’s spectra are derived from population firing rates rather than the LFP, and although oscillations were never a target of its construction or training, the model captured the qualitative depth organization of the mouse V1 SL motif (Fig. 1E, F).

Two key differences should be highlighted. First, the model’s laminar transitions appear sharper than in the recordings, where power smoothly varies with depth. This discrepancy likely arises from the model’s discrete layer structure and, on the other hand, spatial averaging and volume conduction in the data, which tend to blur laminar boundaries in the LFP. Second, the model shows gamma activity more narrowly focused in L4, while experimental maps reveal high-frequency power extending into the supragranular layers. This aligns with our finding that L4 primarily generates 50–80 Hz gamma, which then propagates to L2/3 and L5. The superficial extension observed in recordings likely results from this inter-laminar propagation and the dendritic and volume-conducted components of the LFP that firing-rate spectra do not capture. Overall, the consistency between the model and the experimental maps—covering V1 and various visual areas, and under behavioral tasks distinct from our visual-stimulation protocol—supports the observations from the modeling presented here as an approximate mechanistic explanation of the SL motif, while also delineating where firing-rate proxies and biophysical LFPs are expected to differ.

### L4 is the predominant source of 50–80 Hz gamma in the network

L4 acted as the dominant generator of 50–80 Hz gamma in the model. Silencing L4 strongly suppressed 50–80 Hz gamma across the network, whereas isolated L4 sustained oscillations in this range. Thus, L4 is both intrinsically sufficient to generate the rhythm and is key to its expression across other layers. Importantly, L4 is the strongest but not the only source: isolated L5 retained a reduced 50–80 Hz component, yet L4 clearly dominated and drove stronger oscillations in neighboring layers via inter-laminar excitatory connections. Several features plausibly suit L4 for this role, including its dense recurrent connectivity, high Pvalb density, and strong thalamocortical drive. These results are consistent with experimental observations of granular-layer gamma dominance in mouse V1 ^15,66^, and L4 as the spectral crossover point of the primate SL motif ^7^. Notably, the model’s 50–80 Hz L4 gamma was not inherited from the thalamus, as LGN units showed no corresponding spectral peak (Suppl. Fig. 1C), indicating an intracortical origin distinct from the thalamically driven narrowband gamma reported experimentally ^43–46^.

### Distinct contributions of interneuron subclasses

The perturbation revealed distinct roles for the inhibitory subclasses. Pvalb cells were the principal modulators of the 50–80 Hz gamma; their silencing abolished it network-wide and shifted all layers toward stronger alpha/beta, consistent with their established causal role in fast gamma timing ^16,17,67,68^. Sst cells primarily regulated lower-frequency alpha/beta power, with a more local role supporting L5 gamma, in line with their dendrite-targeting inhibition and reported contributions to gamma maintenance and beta regulation ^16,18^. Vip cells exerted gain control over oscillation magnitude, mainly in superficial layers, without reorganizing frequency structure, consistent with their disinhibitory function ^19^. Thus, the subclasses regulate separable dimensions of the oscillatory state —Pvalb setting fast timing, Sst constraining slower fluctuations, and Vip modulating amplitude through disinhibition—in agreement with models in which distinct interneuron subtypes control distinct features of network synchrony ^25^.

### Temporal properties of Pvalb-to-excitatory synapses are a key determinant of L4 gamma oscillations

Parameter-swapping identified the temporal properties of Pvalb-to-excitatory synapses— specifically the time constants and slow-component amplitude of the double alpha-function—as a key determinant of whether L4 oscillates in the 50–80 Hz gamma regime or the slower alpha/beta regime of L6. Imposing L6-like Pvalb-to-E kinetics on isolated L4 suppressed 50–80 Hz gamma, whereas conferring multiple L4-like properties on L6 induced it. This aligns with the PING framework: fast inhibitory feedback lets the excitatory population recover quickly, permitting faster cycles, whereas slower kinetics lengthen the cycle and favor beta ^53–56^. However, we emphasize that no single parameter or cell type is solely responsible: cell density, connectivity, and intrinsic neuronal properties each contributed, and the full 50–80 Hz rhythm required their convergence. This multifaceted, interdependent nature is a key finding that would be hard for simplified models to predict, highlighting the importance of bio-realistic, multilayer simulations for understanding the mechanisms underlying cortical dynamics and function.

### Limitations of the study

We characterized oscillations using cell-type– and layer-specific population firing-rate spectra rather than the LFP. Although the LFP is more experimentally accessible, firing-rate spectra of individual cell types can be considered more revealing: they directly show the oscillatory activity of the populations of interest and are not affected by remote synaptic or volume-conduction effects that can influence the LFP. On the other hand, population firing-rate spectra do not capture dendritic currents or spatially structured extracellular fields, so features of the experimental LFP that arise from dendritic dipoles—such as contributions of L2/3 and L5 pyramidal apical tufts—to the supragranular signal are not represented, a caveat especially relevant for interpreting L2/3 spectral composition. Additionally, because dendritic-targeted inhibition lacks full compartmental representation in our model, the role of Sst interneurons might be underestimated. However, our point-neuron network model does not produce a biophysical LFP because it lacks the extended transmembrane currents that generate extracellular fields ^29,33,63,69–73^; hence, the population firing rate is both a logical and a practical measure. Previous studies using similar models have also relied on firing-rate spectra (e.g., ^18,19,24–26,35^). This measure and the LFP complement each other, and integrating them in the future will help us compare model predictions more directly with laminar LFP recordings.

Furthermore, we study V1 largely in isolation. In the brain, oscillations are shaped by many factors beyond the local circuit: thalamic inputs carry distinct oscillations ^43–46^, cortico-thalamic interactions via L6 are likely to influence cortical oscillations, and feedback projections impose additional oscillatory modulation ^14,15,74^. Our aim was not to capture all these influences, but to dissect the local cortical mechanisms that likely control both local and brain-wide oscillations. Such mechanisms may, for instance, underlie the proposed prevalence of gamma in feedforward and alpha/beta in feedback processing ^14,15^.

Finally, the model offers a mechanistic hypothesis rather than proof that these synaptic parameters determine the SL motif in vivo; future work should directly compare simulated spikes, forward-modeled LFPs, and cell-type-specific physiological signatures under matched stimuli.

### Conclusions

Together, these results suggest that the laminar organization of cortical rhythms can arise from interactions between sensory drive and layer-and cell-type-specific circuit architecture. In this framework, L4 Pvalb-mediated inhibition is a key (but not only) component that establishes the 50–80 Hz gamma rhythm. Deeper-layer synaptic temporal properties favor slower rhythms, Sst populations regulate low-frequency and lower-gamma oscillations, and Vip populations modulate the balance and gain of oscillatory power. The model thereby provides a mechanistic account of how the SL motif can emerge from cortical circuitry. We have made the model, code, and simulation results publicly available to facilitate future studies of cortical oscillations.

## Methods

### Bio-realistic mouse V1 model

All simulations in this study were performed using the most recent version of the Allen Institute large-scale, bio-realistic model of the mouse primary visual cortex (V1), described in ^20^, which builds directly upon the model of ^21^. Specifically, we used the trained model variant that covers a 400-µm-radius column of V1 and contains approximately 67,000 neurons distributed across the five cortical layers (L1, L2/3, L4, L5, and L6). The model represents 19 distinct cell types: one excitatory (E) and three inhibitory subclasses—parvalbumin (Pvalb)-, somatostatin (Sst)-, and vasoactive intestinal peptide (Vip)-expressing interneurons—per layer for L2/3 through L6, plus a single Htr3a inhibitory subclass in L1. Neurons are modeled as Generalized Leaky Integrate- and-Fire level 3 (GLIF3) point neurons whose intrinsic parameters were individually fit to experimental data from the Allen Cell Types Database, derived from in *vitro* patch-clamp recordings in the mouse cortex ^75,76^.

In the model, synaptic connectivity is grounded in multiple complementary anatomical datasets. Connection probabilities between cell types, as well as the number of synapses per connection, were derived from two large-scale electron-microscopy reconstructions: the V1DD dataset, which provides direct laminar connectivity data from mouse V1 ^77^, and the MICrONS dataset ^22^, used to supplement statistics on in-degree distributions and per-connection synapse counts where V1DD proofreading was less complete. Interneurons in V1DD were assigned to model Pvalb, Sst, or Vip/Htr3a subclasses based on their axonal innervation patterns (proximal-targeting, distal-targeting, or inhibitory-cell-targeting, respectively), and L1 inhibitory somata were assigned to the L1 inhibitory subclass. Synaptic physiology parameters—including postsynaptic potential amplitudes, the time constants of the fast and slow components of the double-alpha synaptic currents, the amplitude of the slow component, and synaptic latencies across cell-type pairs—were obtained from the Allen Institute multipatch synaptic physiology dataset, which comprises systematic paired whole-cell recordings ^23^. Like-to-like connectivity rules reflecting direction-preference similarity were incorporated for excitatory connections (positive like-to-like) and inhibitory connections (negative like-to-like), consistent with the functional organization identified in the MICrONS data.

A key advance of this model generation relative to its predecessor ^21^ is the training procedure, in which synaptic weights were refined using backpropagation through time (BPTT) while keeping the per-cell-type-pair weight distributions consistent with experimental synaptic physiology measurements ^23^. Training improved the match between simulated and in vivo activity— specifically cell-type- and layer-specific firing rates and orientation/direction-selectivity indices— using the Allen Brain Observatory Neuropixels dataset as a target ^27^. Importantly, we did not include oscillatory activity in the construction or training objectives. The resulting trained model exhibits stable spontaneous activity without hand-tuning and achieves substantially improved quantitative agreement with in vivo data compared with the untrained baseline. A complete description of the model and the optimization procedure is given in Ito et al. (2026).

Visual inputs to the model are conveyed via thalamocortical projections from a model of the dorsal lateral geniculate nucleus (LGN). The LGN model generates spike trains whose statistics are determined by the visual stimulus presented to a simulated visual field, with spatial and temporal filter properties derived from published LGN physiology ^21^. LGN afferents project to all cortical layers with layer- and cell-type-specific targeting probabilities consistent with known thalamocortical anatomy. This architecture enables the simulation of arbitrary visual stimuli— including drifting gratings and naturalistic images and movies—all presented as pixel-specific luminance sequences that drive the LGN spike-train generators.

### Visual stimuli

For the main simulations, we presented the LGN model with drifting gratings at a spatial frequency of 0.04 cycles/deg, 0.8 contrast, 2 Hz temporal frequency, and 8 directions (0, 45, 90, 135, 180, 225, 270, and 315°, clockwise from 0° = right-to-left), simulating 10 trials per direction with trial-to-trial randomness in the LGN spike generation. To evaluate the dependence of the laminar oscillation patterns on stimulus parameters, we additionally presented drifting gratings at six contrasts (0.05, 0.1, 0.2, 0.4, 0.6, and 0.8; at 2 Hz, 8 directions) and six temporal frequencies (1, 2, 4, 8, 15, and 30 Hz; at 0.8 contrast, 8 directions), with 10 trials per direction in each condition.

We also presented naturalistic stimuli. The natural scene set comprised 8 sets of 10 natural images each; each image was shown for 250 ms, for a total of 2.5 s per set, following an initial 500 ms gray screen. The natural-movie stimuli consisted of two clips (natural movie one and natural movie two) from the opening scene of *Touch of Evil* (Welles, 1958). Each clip was divided into eight 3 s segments, yielding 8 unique natural-movie input sets; within each segment, frames were presented continuously (12 frames per 250 ms) with no inter-frame interval. For each set, we simulated 10 trials and, unlike for natural scenes, did not present an initial gray screen. The relatively short segment duration was chosen to enable a large number of trials and segments, thereby improving robustness to spectral estimation across stimuli.

### Spontaneous activity

To characterize intrinsic network dynamics in the absence of patterned visual drive, we simulated spontaneous activity by presenting a uniform gray screen, which produces baseline LGN input without stimulus structure. Spontaneous simulations were run with the same number of trials and total duration as the corresponding visually evoked condition, and cell-type– and layer-specific power spectra and SL-motif maps were computed as described below.

### Simulation and power-spectrum analysis

We ran simulations using the Brain Modeling Toolkit (BMTK) ^78^, which interfaces with the NEST simulator, at a temporal resolution of 0.1 ms. For each stimulus condition, network activity was recorded as spike trains from all neurons, from which cell-type– and layer-specific population firing rates were computed (binned at the resolution given below) and used as proxies for the local field potentials (LFPs). We did not normalize the firing rates by the number of neurons per layer; consequently, a layer’s power spectrum magnitude reflects both its firing rates and the number of contributing neurons, which matters when comparing absolute power across layers. This spike-based approach enables tractable, simultaneous characterization of oscillatory dynamics across all cortical layers and cell types, without the additional computational cost of full biophysical LFP modeling.

Power spectra were computed from population firing-rate time series using Welch’s method (segment length nperseg = 500, bin size = 2 ms, scaling = "spectrum"). With the "spectrum" scaling, Welch’s method returns the squared amplitudes of the spectral components rather than a per-Hz spectral density; because the input signal is a firing rate (spikes/s), the resulting power has units of (spikes/s)². We computed spectra at two levels: cell-type-specific spectra from each cell type’s population firing rate within a layer, and layer-specific spectra from the aggregated firing rate of all neurons in that layer. We used a cross-layer (SL motif) normalization to construct the SL motif maps (e.g., Fig. 1D, Fig. 2D, Suppl. Fig. 1B): the power at each frequency was divided by the maximum power across layers at that same frequency, yielding a value between 0 and 1 that indicates which layer dominates at each frequency. Raw, unnormalized spectra are reported in units of (spikes/s)^2^. In Suppl. Fig. 9J, we used within-spectrum (peak) normalization to compare the spectral shape across model variants (e.g., the isolated-layer comparisons in Fig. 6J); each individual power spectrum was divided by its own maximum value.

### Experimentally observed spectrolaminar motif in the mouse visual cortex

To compare the model-predicted SL motif with experimentally observed motifs, we analyzed mouse LFP recordings from two previously published studies ^41,42^. In these studies, researchers recorded neural activity from awake, head-fixed mice using high-density Neuropixels probes spanning cortical depth during visual discrimination ^41^ and oddball ^42^ tasks. The original studies used these data for different analyses and did not report SL-motif maps.

#### Dataset 1 ^41^

From six V1 recordings in four mice, we analyzed sites labeled as VISp. The original authors identified the anatomical locations of the recording sites using histologically reconstructed probe tracks registered to the Allen Common Coordinate Framework. For analysis, we only included sites within the cortical sheet. Cortical depths were normalized to the entire cortical thickness, and estimated layer boundaries were based on the average thickness of individual layers in mouse V1 ^79^. Power spectra were computed with a Hann-tapered fast Fourier transform averaged over 5-s segments. Narrowband electrical noise was removed using a combination of Notch filtering and interpolation of relative power values.

#### Dataset 2 ^42^

We analyzed LFP from six visual cortical areas (V1, LM, RL, AL, PM, AM) in mice. We computed normalized LFP power by cortical layer for all 83 Neuropixels probes across the six visual areas (14 mice, DANDI:000253) over the full trial (the epoch of the grand spectrolaminar analysis: https://github.com/BastosLab/epych/blob/develop/notebooks/passiveglo/grand_spectrolaminar.ipynb: 200 ms windows, six task conditions pooled). The LFP was median-filtered across channels; channels were assigned to layers by their Allen CCF label (L2/3, L4, L5, L6a, L6b; L1 excluded; the CCF does not separate L2 from L3), and every fourth channel was used.

To generate the SL-motif maps for both datasets, we normalized the power at each frequency by the maximum power across depth at that frequency (the same cross-layer/depth SL motif normalization used for the model laminar power), yielding a normalized frequency-by-depth map per recording probe (Suppl. Fig. 1 D). We averaged the SL map across all probes in each dataset to obtain unified SL-motif maps (Fig. 1E-F). Because point-neuron models cannot generate LFP, this comparison between experimental LFP-derived and model firing-rate-derived SL maps is qualitative and focuses on the laminar organization of relative gamma and alpha/beta power rather than absolute power.

### Harmonic relationship assessment

#### Empirical mode decomposition

To test whether the L4 50-80 Hz gamma component was a harmonic of a lower-frequency fundamental oscillation, we analyzed, separately for each of the 10 trials × 8 drifting-grating directions (80 recordings/simulated trials in total), each sampled at 500 Hz (2 ms bins), the isolated L4 population firing rate (while silencing all cell in other layers). Each firing rate time series was decomposed into intrinsic mode functions (IMFs) using empirical mode decomposition (EMD; ^47^ with the EMD Python toolbox (v0.8.1; ^80^). To obtain stable decompositions robust to mode mixing, we used iterated masked EMD (itEMD; ^48^ (2022)), which adds a masking signal during sifting (Deering and Kaiser, 2005) and iteratively re-estimates its frequency until convergence. Sifting was limited to five IMFs, and the converged mask (carrier) frequency of each IMF was retained (in Hz) for every recording. Mask frequencies are summarized per IMF as the median and interquartile range across the 80 simulated trials.

#### Hilbert–Huang transform

The instantaneous phase, frequency, and amplitude of each IMF were obtained from its analytic signal via the Hilbert transform ^47^. Per-IMF amplitude-weighted Hilbert–Huang transforms (HHT) were computed over 128 linearly spaced frequency bins spanning 0–100 Hz. IMF waveforms and HHT spectra were averaged across the 80 recordings and reported as the mean ± standard error of the mean (SEM). Because IMFs are not strictly phase-aligned across recordings, trial-averaged IMF waveforms emphasize the stimulus-locked component; the mean instantaneous amplitude (Hilbert envelope) is also reported as a phase-cancellation-free measure of each mode’s amplitude time course.

#### Harmonic-relationship tests

To determine whether the IMFs reflect independent oscillations rather than harmonics of a common fundamental, we applied the harmonic-criteria framework of ^48^. Taking the slowest mode (IMF-5, ≈2 Hz) as the candidate fundamental, each faster IMF was tested against three criteria, evaluated over 18 approximately equal-length segments: (i) an integer instantaneous-frequency ratio with the fundamental (one-sample t-test of the ratio against its nearest integer; met when p > 0.05); (ii) a consistent phase relationship via phase–phase distance correlation (Székely et al., 2007; met when p < 0.05); and (iii) an amplitude–frequency (af) product below one (one-sided one-sample t-test against 1; met when p < 0.05). We classified an IMF pair as harmonically related only when all three criteria were met simultaneously. We evaluated the criteria per recording and aggregated them across the dataset.

### Network perturbations

#### Cell-type- and layer-specific silencing

To test the role of inhibitory cell types in network oscillations, we silenced all neurons of a given inhibitory subclass (Pvalb, Sst, or Vip)—either across the whole network or layer by layer—by applying a constant hyperpolarizing current throughout the simulation—and simulated the perturbed network’s response to drifting gratings (0.8 contrast, 2 Hz, 8 directions). We then compared the resulting cell-type– and layer-specific power spectra with the unperturbed baseline.

To dissociate the contribution of inter-layer connections from within-layer circuitry, we performed two complementary manipulations under the same visual stimulation. In the **silencing perturbation**, we silenced all neurons in a given layer and examined the effect on the rest of the network. In the **isolated-layer perturbation**, we silenced all neurons except those in a single layer of interest—for example, to isolate L4 we silenced all neurons in L1, L2/3, L5, and L6—and computed the layer-specific power spectrum from the remaining active layer. The same procedure was applied to each layer in turn.

#### Model variants (L4↔L6 parameter swaps)

To identify the circuit properties that generate the 50–80 Hz gamma in L4, we built a series of isolated-L4 model variants in which specific circuit elements were replaced with the corresponding properties of L6—a layer that consistently oscillates near 20 Hz with no 50–80 Hz gamma component across all perturbations tested. Because even small changes to these parameters can strongly affect firing magnitude and network stability, each variant was retrained (as described above, following the approach of ^20^, which) to match the target in vivo firing rates before spectral analysis. We constructed the following variants: L4 with L6 cell volume density; L4 with L6 cell models (intrinsic electrophysiological cell dynamics); L4 with L6 within-layer connectivity; L4 with L6 cell volume density and within-layer connectivity (combined); L4 with L6 synaptic parameters (the fast- and slow-component time constants of the double-alpha synaptic currents, the slow-component amplitude, and the synaptic latency); L4 with L6 E-to-E, E-to-I, and I-to-E synaptic parameters; L4 with L6 Pvalb-to-E, Sst-to-E, and Vip-to-E synaptic parameters (separately). I-to-I connections share the same parameters across layers and were therefore not swapped. For the Pvalb-to-E and Sst-to-E cases, we additionally built sub-variants that swapped (i) the synaptic time constants and alpha-function amplitudes together and (ii) the synaptic latencies alone, to dissociate their contributions.

Conversely, because L6 consistently oscillated at ∼20 Hz with little change across the perturbations, we tested whether L4-like 50–80 Hz gamma could be induced in L6 by transferring L4 properties into the isolated L6 circuit. We built two L6 variants: one incorporating only L4 synaptic parameters, and another incorporating L4 cell volume density, within-layer connectivity, cell models, and synaptic parameters. We then computed the drifting-grating-evoked power spectrum. The modified L6 network differs from L4: it occupies the original L6 volume, contains more neurons (because L6 is substantially thicker), and receives the original L6 external drive.

## Data and software availability

We have made all code available at https://github.com/AllenInstitute/oscillations-biorealistic-mouse-v1-model. Trained model files for all model variants considered and simulations can be downloaded from https://www.dropbox.com/scl/fo/xzcmw1tnkh1kv3kvo4ma4/AAZv3zGDzned--6RMw8bSQQ?rlkey=tr4gkoy6lomha00qiqjrgmrra&st=5ekwfoer&dl=0.

The model was built using the Brain Modeling ToolKit (BMTK) ^78^, employing the SONATA data format ^81^. We trained all model variants using the BMTK/NEST-generated SONATA network files (connectivity and cell-parameter files) as inputs to a TensorFlow-based differentiable simulator ^20^. We implemented all network training in TensorFlow (2.14.0 + CUDA 11.8), using the pipeline described in Ito et al. (2026). All simulations performed in this study were conducted with BMTK FilterNet and PointNet (using NEST simulator 3.6), with the latter using the trained network. We ran training on a high-performance computing cluster using a single NVIDIA A100 GPU (40 GB VRAM) and Intel Xeon Gold 6330N processors.

Mouse V1 LFP recordings from the Steinmetz et al. (2019) dataset^41^, used in Fig. 1E, are publicly available at https://figshare.com/articles/steinmetz/9598406. LFP recordings from the mouse visual cortex in the Sennesh et al. (2025) dataset^42^, used in Fig. 1F, are publicly available through the DANDI Archive (Dandiset ID: 000253; DOI: https://doi.org/10.48324/dandi.000253/0.240503.0152). The corresponding analysis code is available on GitHub: https://github.com/BastosLab/epych/tree/develop/notebooks/passiveglo.

## Supplementary Figures

**Supplementary Figure 1.**
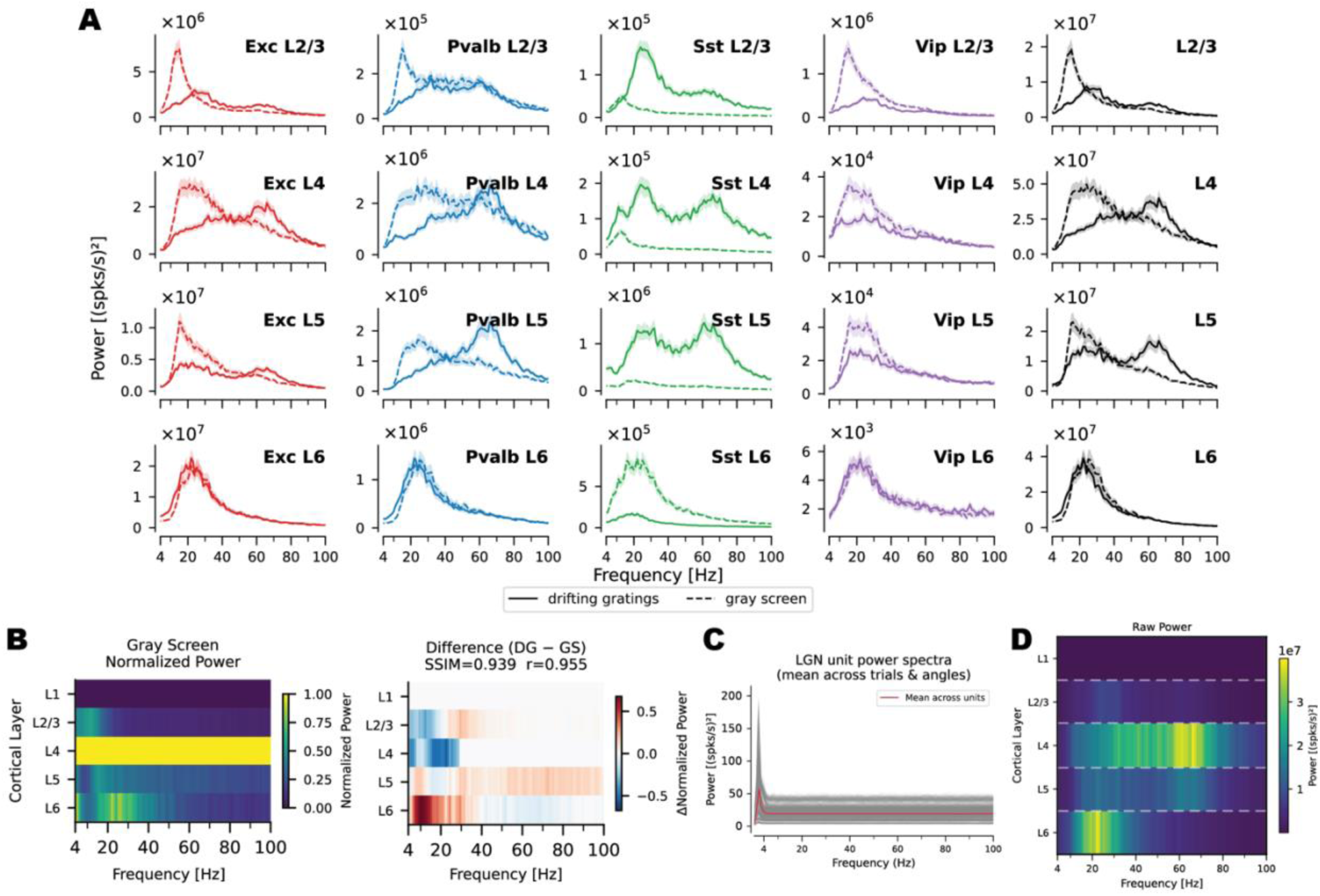
Spontaneous and visually evoked cell-type– and layer-specific power spectra. **(A)** Spontaneous (gray screen) and visually evoked (drifting gratings: 0.8 contrast, 2 Hz temporal frequency) cell-type– and layer-specific power spectra. **(B)** Spontaneous (gray screen; left panel) SL motif map. Normalization was performed relative to the maximum power per frequency across layers. The right panel shows the difference map, highlighting deviations of the visually evoked SL motif (Fig. 1D – right panel) from the spontaneous activity. The structural similarity index (SSIM) and Pearson correlation between the spontaneous SL map and the drifting grating SL map are shown, quantifying the preservation of the SL motif. **(C)** Mean power spectra of LGN units across trials and drifting grating directions (red line). Gray lines show the mean power for each of the 17,400 LGN units. **(D)** V1 model’s raw laminar power spectra revealing the characteristic frequency–layer organization observed in SL motif: gamma-band dominance in L4 and alpha/beta dominance in L6. Dashed lines mark estimated boundaries of cortical layers.

**Supplementary Figure 2.**
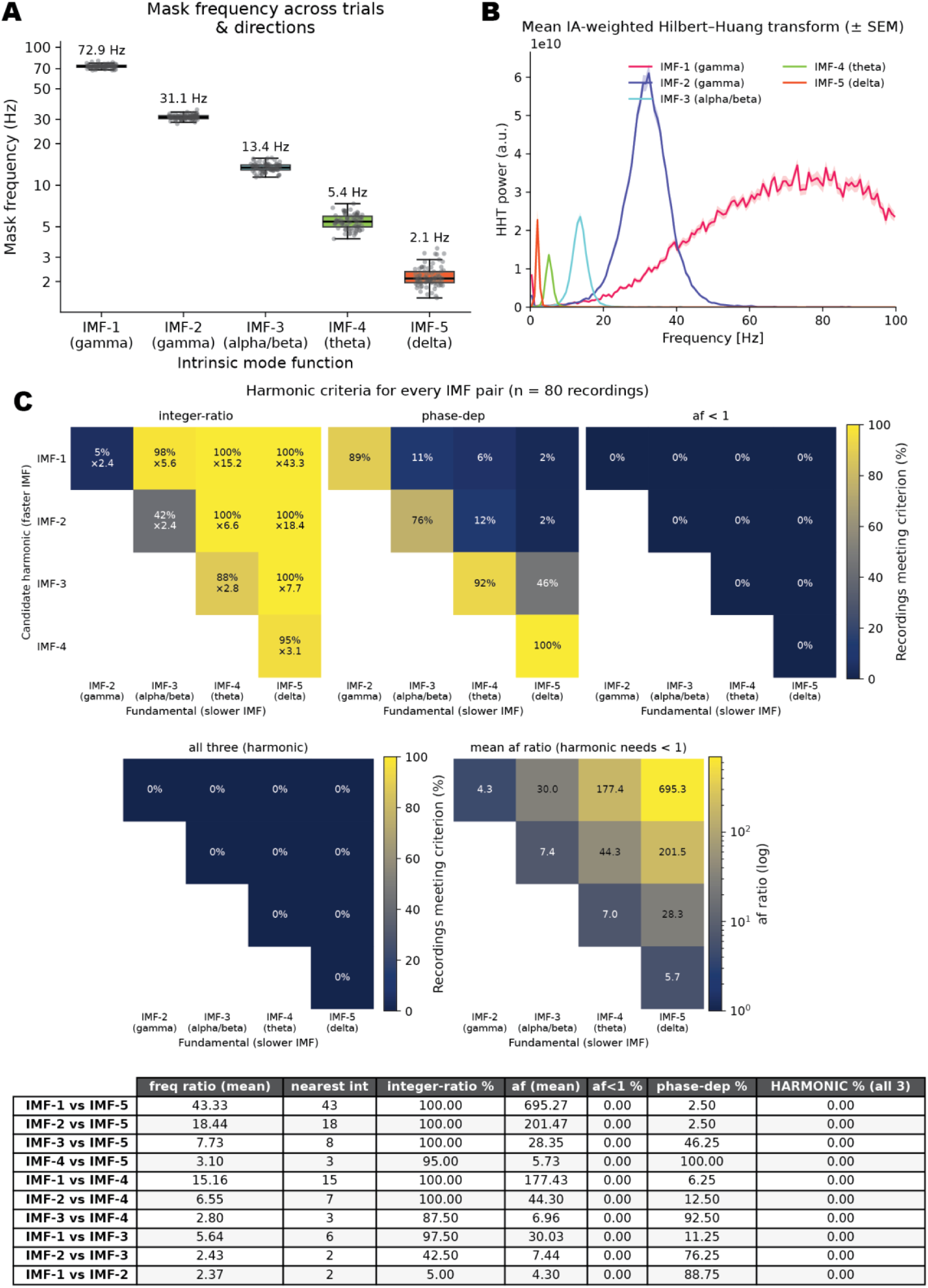
Empirical Mode Decomposition (EMD) of L4 population activity to identify the presence of high-frequency harmonics. **(A)** Distribution of mask (carrier) frequencies for each of the 5 intrinsic mode functions (IMFs) within the L4 population activity across 10 trials and 8 stimulus directions. Gray dots represent individual simulated trials, totaling 80 recordings (10 simulated trials multiplied by 8 directions). **(B)** Amplitude-weighted Hilbert–Huang power spectrum for each IMF, averaged across the 80 recordings, with ± SEM shading. Each IMF occupies a distinct frequency band whose peak matches the corresponding mask frequency in panel A (IMF-5 ≈ 2 Hz delta up to IMF-1 broadband gamma). **(C)** Harmonic-relationship assessments for every IMF pair, following ^48^. Each pairing tests whether a faster IMF (candidate harmonic; matrix rows) is a harmonic of a slower IMF (potential fundamental; matrix columns). All 10 pairings are tested: IMF-1–4 vs IMF-5, IMF-1–3 vs IMF-4, IMF-1–2 vs IMF-3, and IMF-1 vs IMF-2. Three criteria must hold simultaneously for a pairing to be classified as harmonic: (1) the mean instantaneous-frequency ratio does not differ significantly from the nearest integer (p > 0.05; integer-ratio); (2) the phase–phase distance correlation is significant (p < 0.05; phase-dep); and (3) the amplitude–frequency ratio af is significantly below 1 (p < 0.05; af < 1). Matrices: percentage of the 80 recordings meeting each criterion individually and all three jointly ("harmonic"); integer-ratio cells also give the mean frequency ratio (×). The last matrix shows the mean af ratio on a logarithmic color scale. Table: criteria per pairing, averaged across the 80 simulated trials. Adjacent IMFs had frequency ratios of ≈2.4–3.1 and consistent phase coupling (76–100% of recordings), whereas non-adjacent pairings were rarely phase-coupled (≤46%). The two fastest adjacent pairings did not have integer frequency ratios: IMF-1 vs IMF-2 met the criterion in 5% of recordings and IMF-2 vs IMF-3 in 42.5%. All other pairings met it in ≥87.5% of recordings, although this test is weakly constraining for large ratios (≳5). Criterion 3 failed for every pairing in every recording (mean af = 4.3–695.3, well above the af = 1 threshold). No pairing therefore satisfied all three criteria (harmonic = 0% for all 10 pairings; 0 of 800). These results indicate that the five IMFs are independent oscillatory modes rather than harmonics of one another.

**Supplementary Figure 3.**
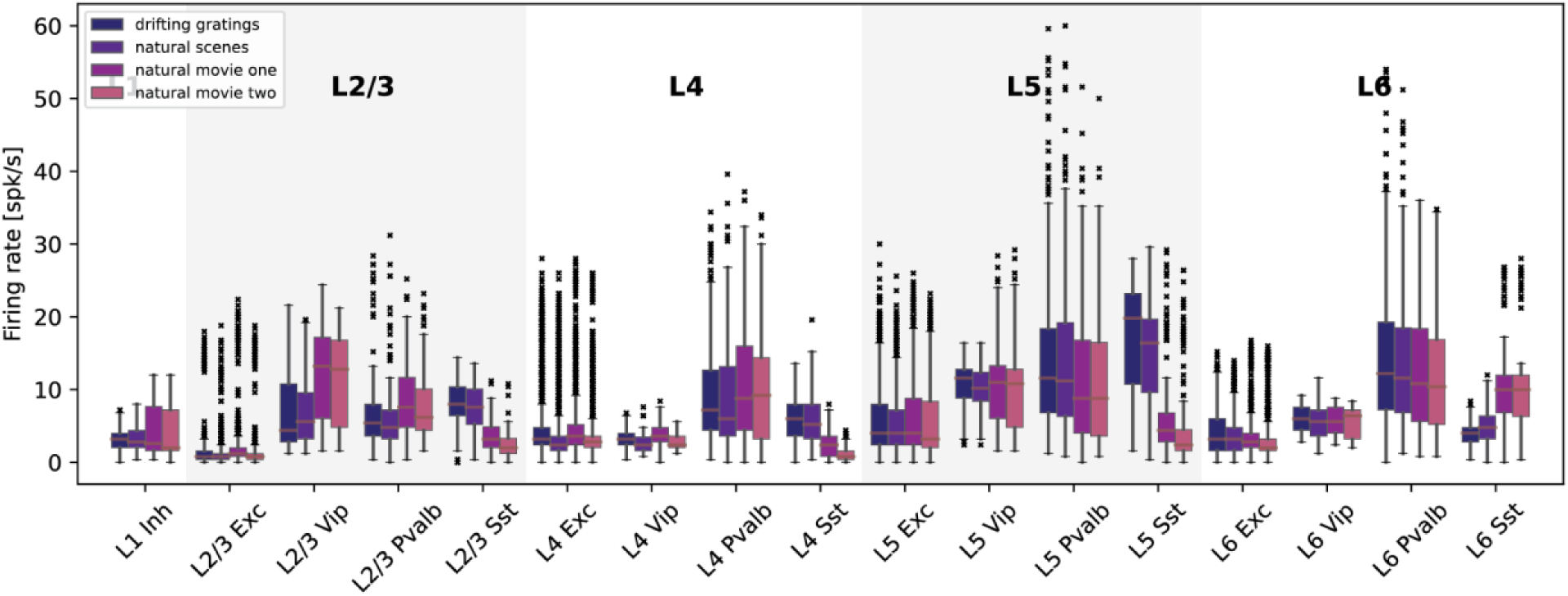
Firing rate distributions across cell types and visual stimulus conditions. Cell-type-specific firing rate distributions for the V1 model in response to drifting gratings (0.8 contrast, 2 Hz temporal frequency), natural scenes, natural movie 1, and natural movie 2. Box plots show the distributions of single-neuron firing rates for a distinct cell type (Exc, Pvalb, Sst, Vip/L1 Inh in a given layer), as labeled, where the firing rate of each neuron is averaged over the duration of the visual stimulus, over stimuli within the group (e.g., all of the 8 directions of the drifting gratings or all of the 8 sets of natural images), and over all trials.

**Supplementary Figure 4.**
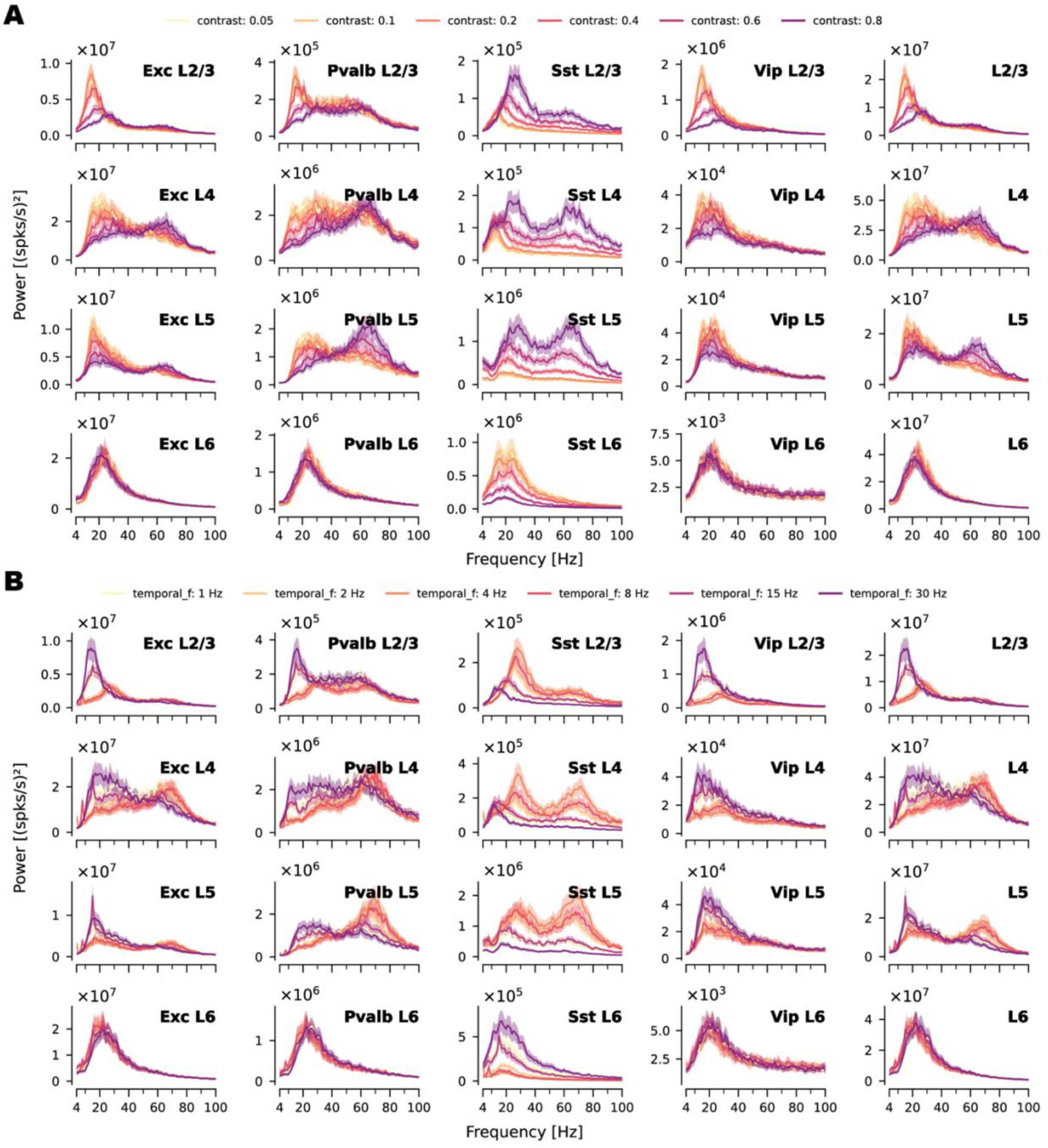
Stimulus contrast and temporal frequency modulate oscillations in a cell-type- and layer-specific manner. **(A)** Cell-type– and layer-specific power spectra evoked by drifting gratings at six contrast levels (0.05, 0.1, 0.2, 0.4, 0.6, and 0.8), at 2 Hz temporal frequency and 8 directions. **(B)** Cell-type– and layer-specific power spectra evoked by drifting gratings at six temporal frequencies (1, 2, 4, 8, 15, and 30 Hz), at 0.8 contrast and 8 directions. Every contrast level and temporal frequency condition had 10 trials per direction.

**Supplementary Figure 5.**
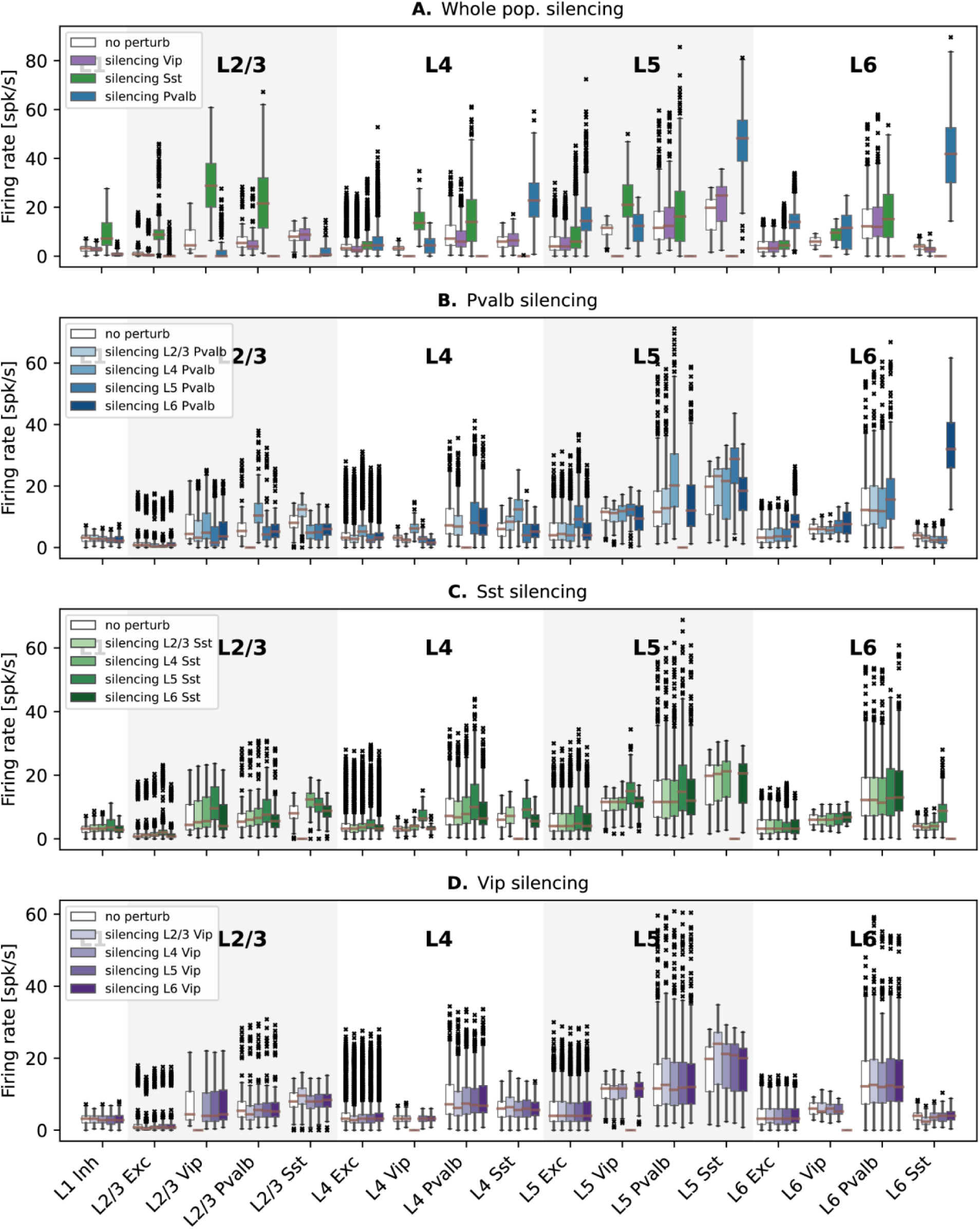
Layer-specific effects of Pvalb, Vip, and Sst silencing on cell-type-specific firing rate distributions. **(A)** Cell-type-specific firing rate distributions for the V1 model in response to drifting gratings (0.8 contrast, 2 Hz temporal frequency) during selective silencing of all Pvalb cells (blue), all Sst cells (green), and all Vip cells (purple). **(B-D)** Cell-type-specific firing rate distributions for the V1 model in response to drifting gratings (0.8 contrast, 2 Hz temporal frequency) during selective layer-by-layer silencing of Pvalb cells (B), Sst cells (C), and Vip cells (D).

**Supplementary Figure 6.**
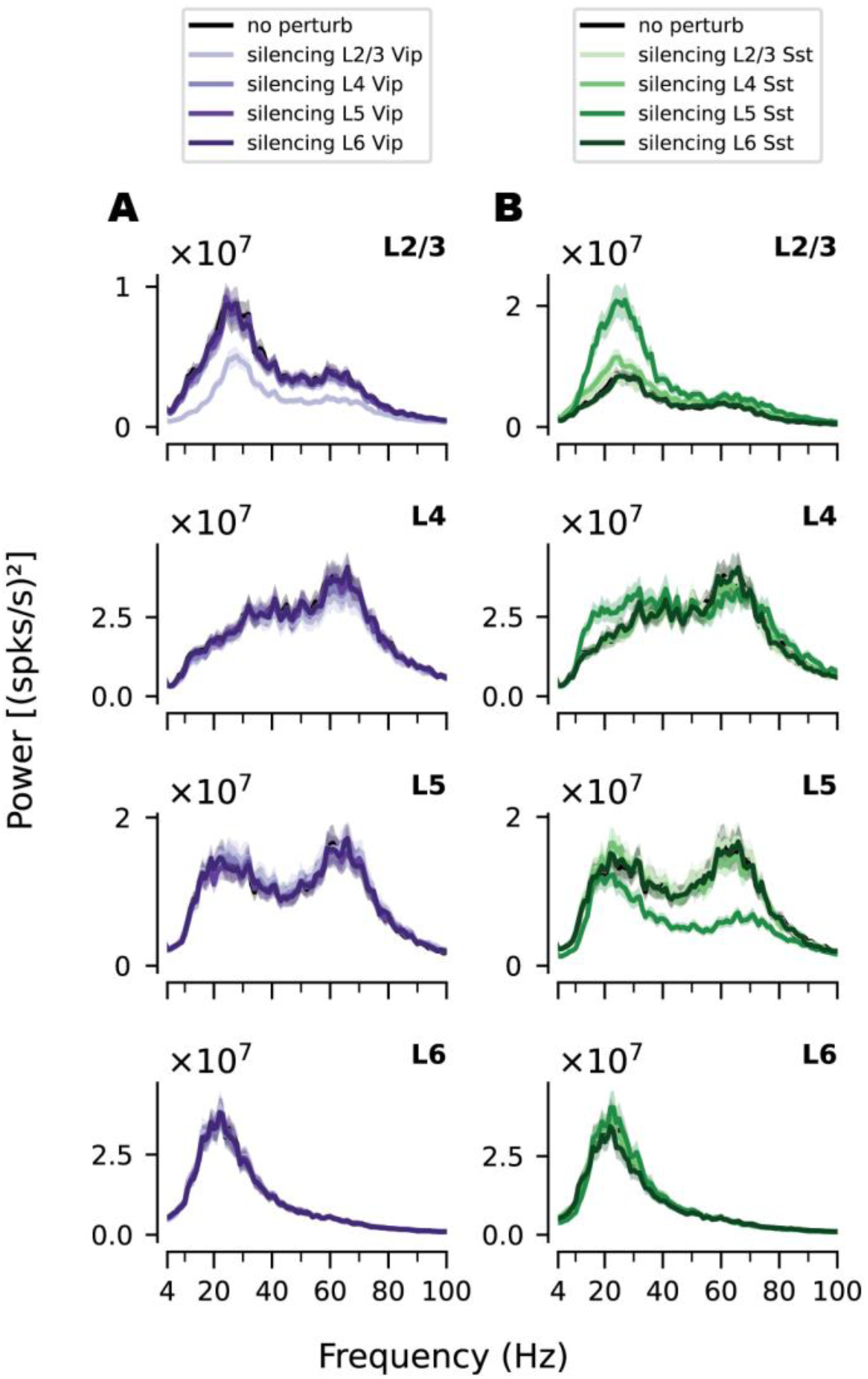
Layer-specific effects of Vip and Sst silencing on oscillatory power. Laminar power spectra following layer-by-layer silencing of Vip **(A)** and Sst **(B)** cells, respectively.

**Supplementary Figure 7.**
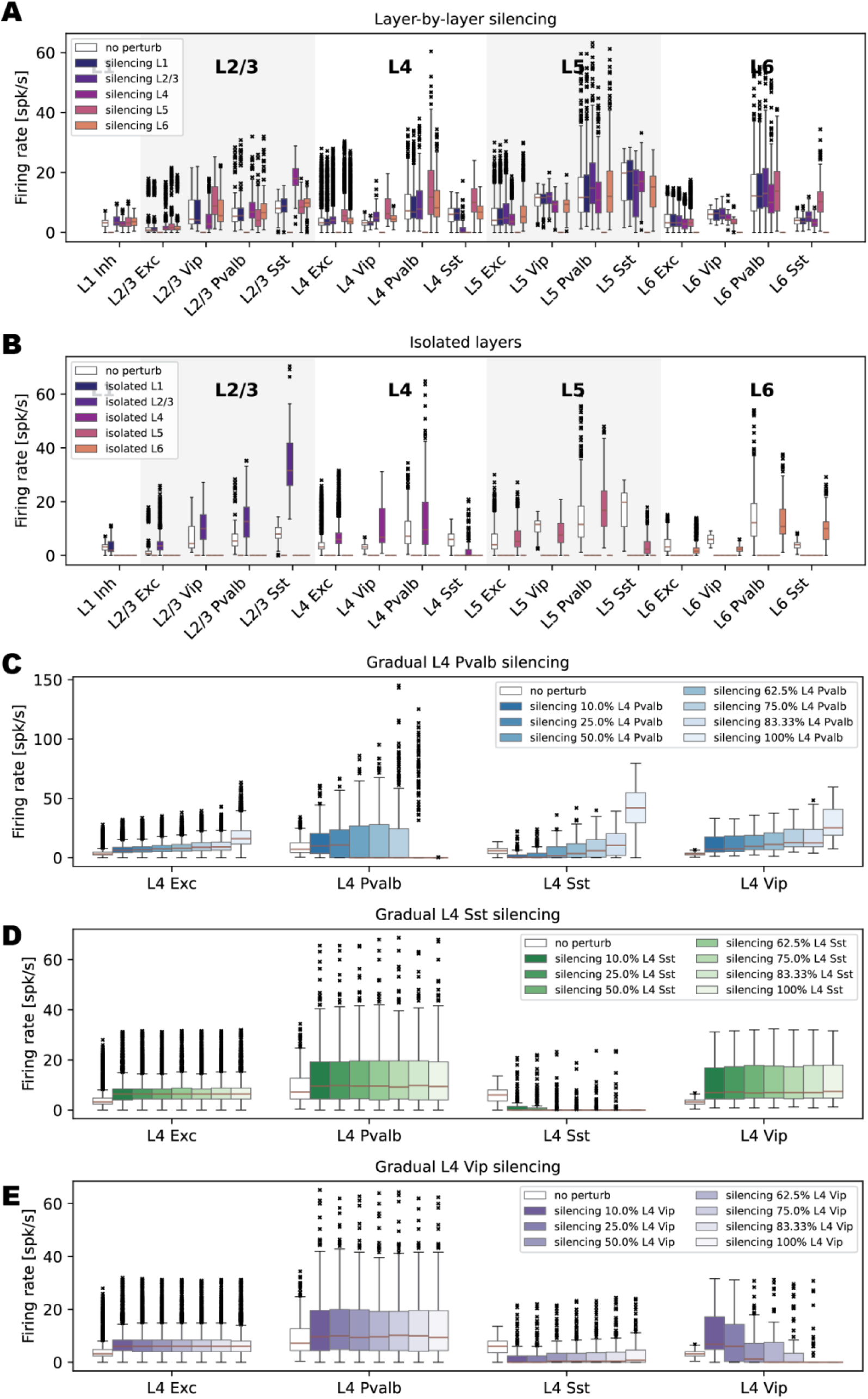
Firing rate distributions across cell types during drifting grating presentation for layer-by-layer silencing and isolated-layer simulations. **(A-B)** Cell-type-specific firing rate distributions for the V1 model in response to drifting gratings (0.8 contrast, 2 Hz temporal frequency) during (A) layer-by-layer silencing and (B) isolated-layer simulations. **(C-D)** Isolated L4 simulation – firing rate distributions of L4 cell types in response to drifting gratings (0.8 contrast, 2 Hz temporal frequency) during gradual silencing of (C) Pvalb, (D) Sst, and (E) Vip cells, respectively.

**Supplementary Figure 8.**
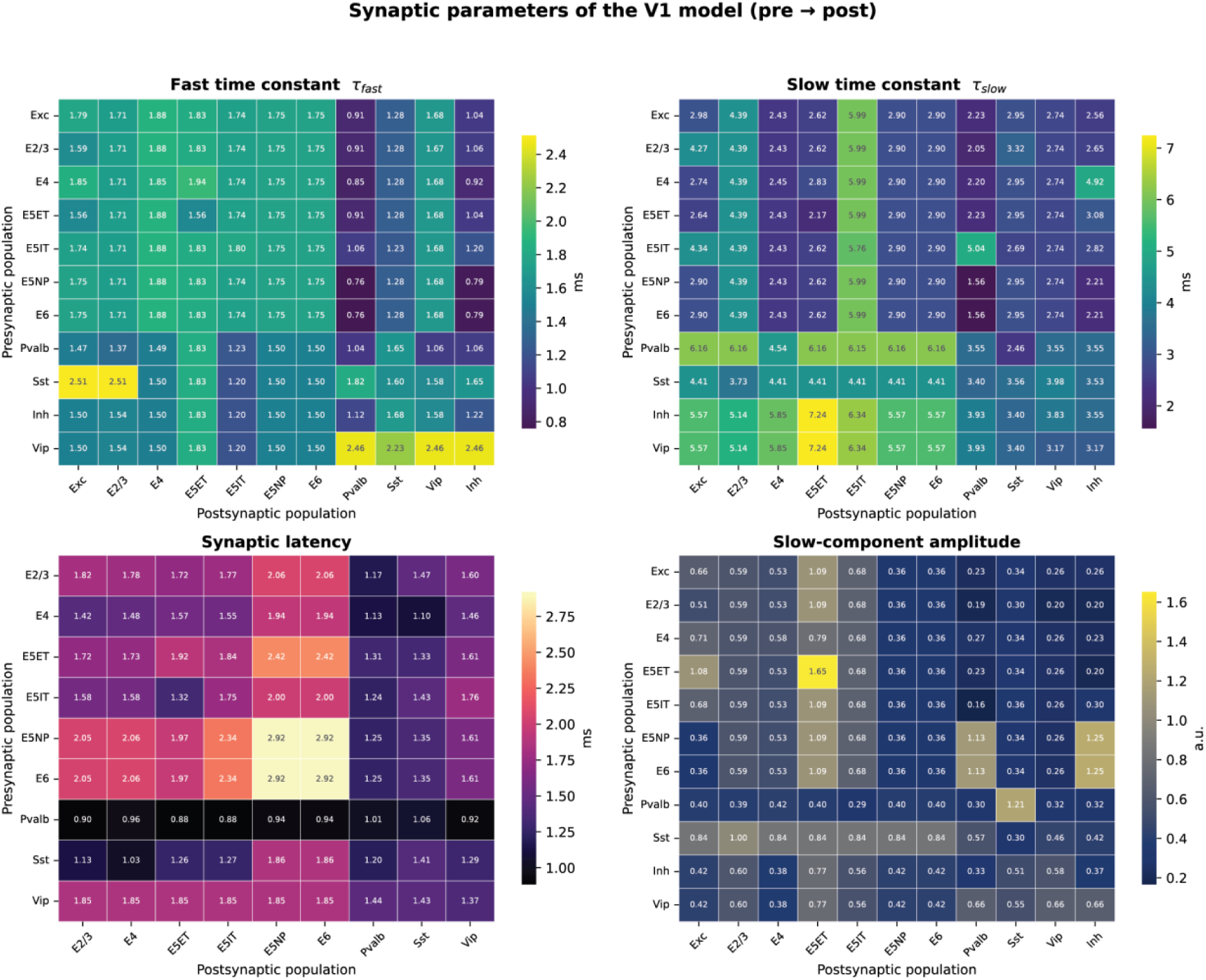
V1 model synaptic parameters (taken from experimental data of ^23^. Top panels – Time constants of the fast (left) and slow (right) components of the double-alpha synaptic currents in the V1 model. Bottom panels – Synaptic latency and amplitude of the slow component of the double-alpha synaptic currents in the V1 model. In all plots, the y-axis indicates the presynaptic populations and the x-axis the postsynaptic population.

**Supplementary Figure 9.**
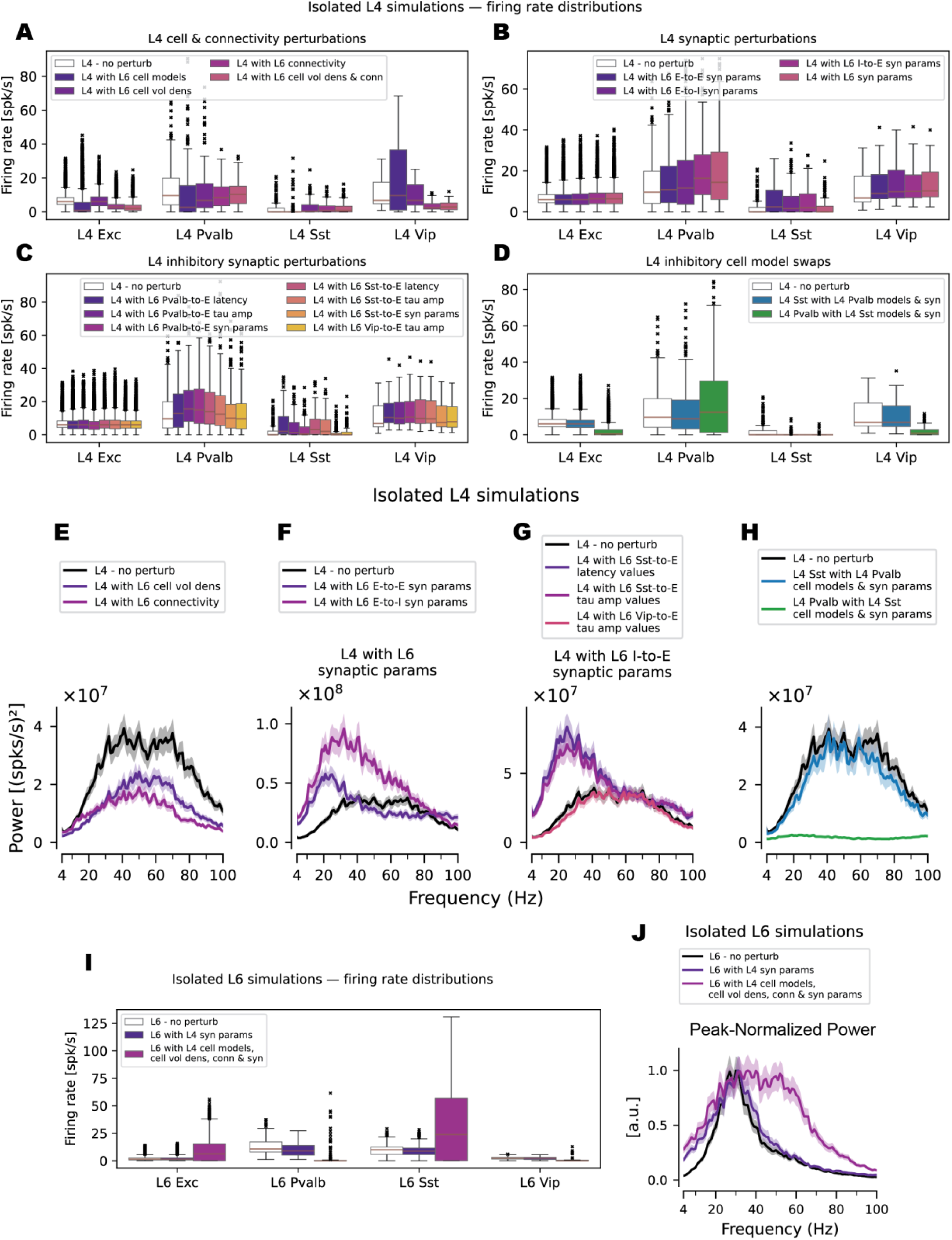
Swapping parameters between L4 and L6. **(A-D)** Cell-type-specific firing rate distributions from simulations of isolated L4 networks, comparing the unperturbed baseline (“L4—no perturbation”) with model variants that systematically replaced L4 network parameters with those from L6. **A:** Variants swapping parameters of cell models (“L4 with L6 cell models”), cell volume density (“L4 with L6 cell vol dens”), within-layer connectivity (“L4 with L6 connectivity”), or cell volume density and within-layer connectivity (“L4 with L6 cell vol dens & conn”). **B:** Variants swapping only excitatory-to-excitatory (E-to-E) synaptic parameters (“L4 with L6 E-to-E syn params”), only excitatory-to-inhibitory (E-to-I) synaptic parameters (“L4 with L6 E-to-I syn params”), only inhibitory-to-excitatory (I-to-E) synaptic parameters (“L4 with L6 I-to-E syn params”), or all synaptic parameters (“L4 with L6 syn params”); **C:** Targeted variants replacing Pvalb-to-excitatory synaptic parameters: all parameters (“L4 with L6 Pvalb-to-E syn params”), only synaptic latency (“L4 with L6 Pvalb-to-E latency values”), or only time constants and alpha-function amplitude (“L4 with L6 Pvalb-to-E tau amp values”); the Sst-to-excitatory synaptic parameters: all parameters (“L4 with L6 Sst-to-E syn params”), only synaptic latency (“L4 with L6 Sst-to-E latency values”), or only time constants and alpha-function amplitude (“L4 with L6 Sst-to-E tau amp values”); and the Vip-to-excitatory counterpart (“L4 with L6 Vip-to-E tau amp values”). **D:** Variants swapping L4 Sst synaptic parameters and cell model parameters with those of L4 Pvalb cells (“L4 Sst with L4 Pvalb cell models & syn params”), and vice versa (“L4 Pvalb with L4 Sst cell models & syn params”). **(E-H)** Power spectra from simulations of isolated L4 networks, comparing the unperturbed baseline (“L4—no perturbation”) with model variants that systematically replaced L4 network parameters with those from L6. **E:** Variants swapping cell volume density (“L4 with L6 cell vol dens”) or within-layer connectivity (“L4 with L6 connectivity”). **F:** Variants swapping only excitatory-to-excitatory (E-to-E) or excitatory-to-inhibitory (E-to-I) synaptic parameters (“L4 with L6 E-to-E syn params” and “L4 with L6 E-to-E syn params”, respectively). **G:** Targeted variants replacing Sst-to-excitatory synaptic parameters—only synaptic latency (“L4 with L6 Sst-to-E latency values”), or only time constants and alpha-function amplitude (“L4 with L6 Sst-to-E tau amp values”)—and the Vip-to-excitatory parameters (“L4 with L6 Vip-to-E tau amp values”). Vip-to-E latency values were the same across layers; thus, only “L4 with L6 Vip-to-E tau amp values” variant was considered. **H:** Variants swapping L4 Sst cells’ synaptic parameters and intrinsic electrophysiological parameters with those from L4 Pvalb cells (“L4 Sst with L4 Pvalb cell models & syn params”), and vice versa (“L4 Pvalb with L4 Sst cell models & syn params”). **(I)** Cell-type-specific firing rate distributions from simulations of isolated L6 networks, comparing the unperturbed L6 baseline with variants in which L6 parameters were replaced with those from L4: L6 with L4 synaptic parameters alone (“L6 with L4 syn params”) and L6 using all internal parameters from L4 (i.e, cell models, cell volume density, connectivity, and synaptic parameters are borrowed from L4; the volume occupied by the network and external inputs remain the same as in the original L6 network). **(J)** Power spectra normalized by their maxima (peak-normalized power) for simulations of the isolated L6 network, comparing the unperturbed L6 baseline with variants in which L6 network parameters were replaced with those from L4: L6 using L4 synaptic parameters alone (“L6 with L4 syn params”) and L6 using all internal parameters from L4.

## References

1. Buzsáki, G. & Vöröslakos, M. Brain rhythms have come of age. Neuron 111, 922–926 (2023).

2. Buzsáki, G. Rhythms of the Brain. (Oxford University Press, 2006). doi:10.1093/acprof:oso/9780195301069.001.0001.

3. Buzsáki, G. & Draguhn, A. Neuronal olscillations in cortical networks. Science 304, 1926– 1929 (2004).

4. Fries, P. Rhythms for Cognition: Communication through Coherence. Neuron 88, 220–235 (2015).

5. Fries, P. A mechanism for cognitive dynamics: neuronal communication through neuronal coherence. Trends Cogn. Sci. 9, 474–480 (2005).

6. Fries, P. Neuronal gamma-band synchronization as a fundamental process in cortical computation. Annual Review of Neuroscience 32, 209–224 (2009).

7. Mendoza-Halliday, D. et al. A ubiquitous spectrolaminar motif of local field potential power across the primate cortex. Nat. Neurosci. 27, 547–560 (2024).

8. Mackey, C. A. et al. Is there a ubiquitous spectrolaminar motif of local field potential power across primate neocortex? bioRxiv 2024.09.18.613490 (2024) doi:10.1101/2024.09.18.613490.

9. Major, A. J. et al. A ubiquitous spectrolaminar motif across independent studies, including in the Mackey et al. dataset. https://doi.org/10.1101/2025.05.07.652644 (2025) doi:10.1101/2025.05.07.652644.

10. Carlson, B. M. et al. V1 interlaminar coherence decreases with interocular conflict. bioRxiv 2025.10.04.680461 (2025) doi:10.1101/2025.10.04.680461.

11. Garcia-Reyes, R., Vezoli, J. & Valdes–Sosa, P. A. Spatial transcriptomic programs relate to spectrolaminar rhythms across macaque cortex. https://doi.org/10.64898/2026.07.08.737187 (2026) doi:10.64898/2026.07.08.737187.

12. Lichtenfeld, M. J. et al. The laminar organization of cell types in macaque cortex and its relationship to neuronal oscillations. bioRxiv 2024.03.27.587084 (2024) doi:10.1101/2024.03.27.587084.

13. Wang, T., Lee, E. K., Carr, N., Li, Y. & Chandrasekaran, C. Distinct neural dynamics in prefrontal and premotor cortex during flexible perceptual decisions. bioRxiv 2026.02.02.702013 (2026) doi:10.64898/2026.02.02.702013.

14. Bastos, A. M. et al. Visual Areas Exert Feedforward and Feedback Influences through Distinct Frequency Channels. Neuron 85, 390–401 (2015).

15. Kerkoerle, T. van et al. Alpha and gamma oscillations characterize feedback and feedforward processing in monkey visual cortex. Proc. Natl. Acad. Sci. 111, 14332–14341 (2014).

16. Chen, G. et al. Distinct Inhibitory Circuits Orchestrate Cortical beta and gamma Band Oscillations. Neuron 96, 1403–1418.e6 (2017).

17. Onorato, I. et al. Distinct roles of PV and Sst interneurons in visually induced gamma oscillations. Cell Rep. 44, 115385 (2025).

18. Veit, J., Hakim, R., Jadi, M. P., Sejnowski, T. J. & Adesnik, H. Cortical gamma band synchronization through somatostatin interneurons. Nature Neuroscience 20, 951–959 (2017).

19. Veit, J., Handy, G., Mossing, D. P., Doiron, B. & Adesnik, H. Cortical VIP neurons locally control the gain but globally control the coherence of gamma band rhythms. Neuron 0, (2022).

20. Ito, S. et al. Deep-learning-assisted simulation of a cortical circuit: integrating anatomy, physiology and function. bioRxiv 2026.03.13.711751 (2026) doi:10.64898/2026.03.13.711751.

21. Billeh, Y. N. et al. Systematic Integration of Structural and Functional Data into Multi-scale Models of Mouse Primary Visual Cortex. Neuron 106, 388–403.e18 (2020).

22. Consortium, T. Mic., et al. Functional connectomics spanning multiple areas of mouse visual cortex. Nature 640, 435–447 (2025).

23. Campagnola, L. et al. Local connectivity and synaptic dynamics in mouse and human neocortex. Science 375, (2022).

24. Tahvili, F., Vinck, M. & Volo, M. di. A mean-field model of neural networks with PV and SOM interneurons reveals connectivity-based mechanisms of gamma oscillations. bioRxiv 2025.10.23.684108 (2025) doi:10.1101/2025.10.23.684108.

25. Tahvili, F., Vinck, M. & Volo, M. di. PV and SOM cells play distinct causal roles in controlling network oscillations and stability. Cell Rep. 44, 116131 (2025).

26. Milea, D., Meneghetti, N., Mazzoni, A. & Cataldo, E. A spiking LIF model captures the role of Somatostatin and Parvalbumin neurons in generating oscillations in V1. bioRxiv 2025.01.24.634718 (2025) doi:10.1101/2025.01.24.634718.

27. Siegle, J. H. et al. Survey of spiking in the mouse visual system reveals functional hierarchy. Nature 592, 86–92 (2021).

28. Mazzoni, A. et al. Computing the Local Field Potential (LFP) from Integrate-and-Fire Network Models. PLoS Computational Biology 11, e1004584 (2015).

29. Hagen, E. et al. Hybrid scheme for modeling local field potentials from point-neuron networks. Cerebral Cortex 26, 4461–4496 (2016).

30. Meneghetti, N., Rimehaug, A. E., Einevoll, G. T., Mazzoni, A. & Ness, T. V. Kernel-based LFP estimation in detailed large-scale spiking network model of mouse visual cortex. Neural Netw. 107866 (2025) doi:10.1016/j.neunet.2025.107866.

31. Nunez, P. L. & Srinivasan, R. Electric Fields of the Brain. (Oxford University Press, 2006). doi:10.1093/acprof:oso/9780195050387.001.0001.

32. Halnes, G., et al. Electric Brain Signals: Foundations and Applications of Biophysical Modeling. (Cambridge University Press, Cambridge, 2024). doi:10.1017/9781009039826.

33. Herrera, B., Sajad, A., Woodman, G. F., Schall, J. D. & Riera, J. J. A Minimal Biophysical Model of Neocortical Pyramidal Cells: Implications for Frontal Cortex Microcircuitry and Field Potential Generation. J. Neurosci. 40, 8513–8529 (2020).

34. Herrera, B., Sajad, A., Errington, S. P., Schall, J. D. & Riera, J. J. Cortical origin of theta error signals. Cereb. Cortex 33, 11300–11319 (2023).

35. Moreni, G., Zou, L., Pennartz, C. M. A. & Mejias, J. F. Synaptic plasticity facilitates oscillations in a V1 cortical column model with multiple interneuron types. Front. Comput. Neurosci. 19, 1568143 (2025).

36. Ponzi, A., Dura-Bernal, S. & Migliore, M. Theta-gamma phase amplitude coupling in a hippocampal CA1 microcircuit. PLOS Computational Biology 19, e1010942 (2023).

37. Henrie, J. A. & Shapley, R. LFP Power Spectra in V1 Cortex: The Graded Effect of Stimulus Contrast. J. Neurophysiol. 94, 479–490 (2005).

38. Perrenoud, Q., Pennartz, C. M. A. & Gentet, L. J. Membrane Potential Dynamics of Spontaneous and Visually Evoked Gamma Activity in V1 of Awake Mice. PLoS Biol. 14, e1002383 (2016).

39. Vries, S. E. J. de et al. A large-scale standardized physiological survey reveals functional organization of the mouse visual cortex. Nat. Neurosci. 23, 138–151 (2020).

40. Schneider-Mizell, C. M. et al. Inhibitory specificity from a connectomic census of mouse visual cortex. Nature 640, 448–458 (2025).

41. Steinmetz, N. A., Zatka-Haas, P., Carandini, M. & Harris, K. D. Distributed coding of choice, action and engagement across the mouse brain. Nature 576, 266–273 (2019).

42. Sennesh, E., Westerberg, J. A., Spencer-Smith, J. & Bastos, A. Ubiquitous predictive processing in the spectral domain of sensory cortex. https://doi.org/10.7554/elife.109053.1 (2025) doi:10.7554/elife.109053.1.

43. Shin, D., Peelman, K., Lien, A. D., Rosario, J. D. & Haider, B. Narrowband gamma oscillations propagate and synchronize throughout the mouse thalamocortical visual system. Neuron 111, 1076–1085.e8 (2023).

44. Meyerolbersleben, L. S., Sirota, A. & Busse, L. Anatomically resolved oscillatory bursts reveal dynamic motifs of thalamocortical activity during naturalistic stimulus viewing. Neuron 113, 2196–2214.e6 (2025).

45. Ramanathan, C. & Veit, J. Lights, contrast, action! Disentangling visually induced oscillations. Neuron 113, 2031–2033 (2025).

46. Saleem, A. B. et al. Subcortical Source and Modulation of the Narrowband Gamma Oscillation in Mouse Visual Cortex. Neuron 93, 315–322 (2017).

47. Huang, N. E. et al. The empirical mode decomposition and the Hilbert spectrum for nonlinear and non-stationary time series analysis. *Proc. R. Soc. Lond. Ser. A: Math., Phys*. Eng. Sci. 454, 903–995 (1998).

48. Fabus, M. S., Woolrich, M. W., Warnaby, C. W. & Quinn, A. J. Understanding Harmonic Structures Through Instantaneous Frequency. *IEEE Open J*. Signal Process. 3, 320–334 (2022).

49. Millman, D. J. et al. VIP interneurons in mouse primary visual cortex selectively enhance responses to weak but specific stimuli. eLife 9, e55130 (2020).

50. Cammarata, C. M. et al. Behavioral state and stimulus strength regulate the role of somatostatin interneurons in stabilizing network activity. Cell Rep. 44, 115954 (2025).

51. Pfeffer, C. K., Xue, M., He, M., Huang, Z. J. & Scanziani, M. Inhibition of inhibition in visual cortex: the logic of connections between molecularly distinct interneurons. Nat. Neurosci. 16, 1068–1076 (2013).

52. Tremblay, R., Lee, S. & Rudy, B. GABAergic Interneurons in the Neocortex: From Cellular Properties to Circuits. Neuron 91, 260–292 (2016).

53. Tiesinga, P. & Sejnowski, T. J. Cortical Enlightenment: Are Attentional Gamma Oscillations Driven by ING or PING? Neuron 63, 727–732 (2009).

54. Wang, X.-J. & Buzsáki, G. Gamma Oscillation by Synaptic Inhibition in a Hippocampal Interneuronal Network Model. The Journal of Neuroscience 16, 6402–6413 (1996).

55. Buzsáki, G. & Wang, X.-J. Mechanisms of Gamma Oscillations. Annual Review of Neuroscience 35, 203–225 (2012).

56. Wang, X.-J. Neurophysiological and Computational Principles of Cortical Rhythms in Cognition. Physiological Reviews 90, 1195–1268 (2010).

57. Börgers, C., Epstein, S. & Kopell, N. J. Gamma oscillations mediate stimulus competition and attentional selection in a cortical network model. Proc. Natl. Acad. Sci. 105, 18023–18028 (2008).

58. Doiron, B., Longtin, A., Turner, R. W. & Maler, L. Model of Gamma Frequency Burst Discharge Generated by Conditional Backpropagation. J. Neurophysiol. 86, 1523–1545 (2001).

59. Masuda, N. & Doiron, B. Gamma Oscillations of Spiking Neural Populations Enhance Signal Discrimination. PLoS Comput. Biol. 3, e236 (2007).

60. Parker, M. M., Rubin, J. E. & Huang, C. State modulation in spatial networks with three interneuron subtypes. Sci. Adv. 11, eads9134 (2025).

61. Bos, H., Diesmann, M. & Helias, M. Identifying Anatomical Origins of Coexisting Oscillations in the Cortical Microcircuit. PLoS Comput. Biol. 12, e1005132 (2016).

62. Dura-Bernal, S. et al. Multiscale model of primary motor cortex circuits predicts in vivo cell-type-specific, behavioral state-dependent dynamics. Cell Reports 42, 112574 (2023).

63. Dura-Bernal, S. et al. Large-Scale Mechanistic Models of Brain Circuits with Biophysically and Morphologically Detailed Neurons. J. Neurosci. 44, e1236242024 (2024).

64. Rimehaug, A. E. et al. Uncovering circuit mechanisms of current sinks and sources with biophysical simulations of primary visual cortex. eLife 12, 2022.02.22.481540 (2023).

65. Rimehaug, A. E., Dale, A. M., Arkhipov, A. & Einevoll, G. T. Uncovering population contributions to the extracellular potential in the mouse visual system using Laminar Population Analysis. PLOS Comput. Biol. 20, e1011830 (2024).

66. Senzai, Y., Fernandez-Ruiz, A. & Buzsáki, G. Layer-Specific Physiological Features and Interlaminar Interactions in the Primary Visual Cortex of the Mouse. Neuron 101, 500–513.e5 (2019).

67. Cardin, J. A. et al. Driving fast-spiking cells induces gamma rhythm and controls sensory responses. Nature 459, 663–667 (2009).

68. Sohal, V. S., Zhang, F., Yizhar, O. & Deisseroth, K. Parvalbumin neurons and gamma rhythms enhance cortical circuit performance. Nature 459, 698–702 (2009).

69. Herrera, B. et al. Resolving the mesoscopic missing link: Biophysical modeling of EEG from cortical columns in primates. NeuroImage 263, 119593 (2022).

70. Næss, S. et al. Biophysically detailed forward modeling of the neural origin of EEG and MEG signals. NeuroImage 225, 117467 (2021).

71. Hagen, E. et al. Brain signal predictions from multi-scale networks using a linearized framework. PLOS Computational Biology 18, e1010353 (2022).

72. Einevoll, G. T., Kayser, C., Logothetis, N. K. & Panzeri, S. Modelling and analysis of local field potentials for studying the function of cortical circuits. Nature Reviews Neuroscience 14, 770–785 (2013).

73. Nunez, P. L. & Srinivasan, R. Electric Fields of the Brain. 203–243 (2006) doi:10.1093/acprof:oso/9780195050387.003.0005.

74. Michalareas, G. et al. Alpha-Beta and Gamma Rhythms Subserve Feedback and Feedforward Influences among Human Visual Cortical Areas. Neuron 89, 384–397 (2016).

75. Teeter, C. et al. Generalized leaky integrate-and-fire models classify multiple neuron types. Nature Communications 9, (2018).

76. Gouwens, N. W. et al. Classification of electrophysiological and morphological neuron types in the mouse visual cortex. Nature Neuroscience 22, 1182–1195 (2019).

77. Institute, A. V1 Deep Dive dataset. https://github.com/AllenInstitute/v1dd_physiology (2024).

78. Dai, K. et al. Brain Modeling ToolKit: An open source software suite for multiscale modeling of brain circuits. PLOS Computational Biology 16, e1008386 (2020).

79. Hage, T. A. et al. Synaptic connectivity to L2/3 of primary visual cortex measured by two-photon optogenetic stimulation. eLife 11, e71103 (2022).

80. Quinn, A. J. et al. Within-cycle instantaneous frequency profiles report oscillatory waveform dynamics. J. Neurophysiol. 126, 1190–1208 (2021).

81. Dai, K. et al. The SONATA data format for efficient description of large-scale network models. PLoS Computational Biology 16, e1007696 (2020).

